# Insights into ancestry and adaptive evolution of the *Mycobacterium tuberculosis* complex from analysis of the emerging pathogen *Mycobacterium riyadhense*

**DOI:** 10.1101/728923

**Authors:** Qingtian Guan, Musa Garbati, Sara Mfarrej, Talal AlMutairi, Alicia Smyth, Albel Singh, Shamsudeen Fagbo, John A. Browne, Muhammad Amin urRahman, Alya Alruwaili, Anwar Hoosen, Conor J Meehan, Chie Nakajima, Yasuhiko Suzuki, Apoorva Bhatt, Stephen V. Gordon, Faisal AlAsmari, A. Pain

**Affiliations:** Pathogen Genomics Laboratory, BESE Division, King Abdullah University of Science and Technology (KAUST), Thuwal-Jeddah, Kingdom of Saudi Arabia; King Fahad Medical City (KFMC), Riyadh, Saudi Arabia; UCD School of Veterinary Medicine, University College Dublin, Dublin, D04 W6F6, Ireland; Institute of Microbiology and Infection, School of Biosciences, University of Birmingham, Edgbaston, Birmingham, UK; One Health Unit, Executive Directorate for Surveillance and Response, Saudi Center for Disease Prevention and Control, Riyadh, Saudi Arabia; School of Chemistry and Biosciences, University of Bradford, Bradford, UK; Global Institution for Collaborative Research and Education, Hokkaido University, Kita 20 Nishi 10, Kita-ku, Sapporo, Japan; Research Center for Zoonosis Control, Hokkaido University, Kita 20 Nishi 10, Kita-ku, Sapporo, Japan

## Abstract

Current evolutionary scenarios posit the emergence of *Mycobacterium tuberculosis*, the deadliest bacterial pathogen for humans globally, from an environmental saprophyte through a cumulative process of genome adaptation. *Mycobacterium riyadhense* is a novel non-tuberculous mycobacterium (NTM) that is being increasingly isolated from human clinical cases with tuberculosis (TB)-like symptoms in various parts of the world. We provide evidence here that *M. riyadhense* is likely a ‘missing link’ in our understanding of the evolution of *M. tuberculosis*. To elucidate the genomic hallmarks that define the evolutionary relationship between *M. riyadhense* and other mycobacterial species, including members of the *Mycobacterium tuberculosis* complex (MTBC), eight clinical isolates of *M. riyadhense* were sequenced and analyzed. We show, among other features, that *M. riyadhense* shares a large number of conserved orthologues with the MTBC; contains linear and circular plasmids carrying type IV and type VII secretion systems; and shows expansion of toxin/anti-toxin pairs. We conclude that *M. riyadhense* is an emerging mycobacterial pathogen that shares a common ancestor with members of the MTBC and that can serve as an experimental model to study the evolution and pathogenesis of tubercle bacilli.

**Author summary:** *Mycobacterium tuberculosis* is one of the most prolific infectious killers in humans and is a member of the *Mycobacterium tuberculosis* complex (MTBC) - a group of genetically related pathogens that cause tuberculosis (TB) in mammalian species. It is postulated that MTBC has evolved from a free-living environmental ancestor into an obligate pathogen. In this evolutionary context, a comprehensive understanding of the genomic hallmarks of the free-living environmental ancestors of the MTBC is of particular scientific interest for better understanding of the evolution of the MTBC. *Mycobacterium riyadhense* is a novel environmental mycobacterium, first isolated in 2009 in a hospital in Riyadh, that is increasingly being isolated from clinical cases with typical tuberculosis (TB)-like symptoms in humans. In this study, we report the characterization of eight clinical isolates of *M. riyadhense*, compare their genomes to members of the MTBC, and provide a comprehensive insight into the adaptive changes associated with the evolution of the MTBC from environmental mycobacteria. We show that *M. riyadhense* is one of the closest known environmental mycobacteria related to the MTBC, and we provide several lines of molecular evidence that *M. riyadhense* is likely the ‘missing link’ in the evolution of *M. tuberculosis*. It shares a common ancestor with members of the MTBC that have evolved through a process of genome reduction, expansion of toxin/antitoxin (T/A) gene systems, and ultimately host adaptation.

## Introduction

The *Mycobacterium tuberculosis* complex (MTBC) is a group of genetically related pathogens that cause tuberculosis (TB) in mammalian species. The hallmark member, *Mycobacterium tuberculosis*, is the single most deadly pathogen, causing over 1.6 million deaths globally in 2017. Current evolutionary scenarios posit the evolution of the MTBC from an environmental saprophyte through a cumulative process of genome adaptation. Such scenarios envisage intermediate mycobacterial species with increasing pathogenic potential for humans, the vestiges of which should be present in extant mycobacterial species. Comparative genomic analyses between the MTBC members and opportunistic mycobacterial pathogens may therefore reveal the key evolutionary steps involved in the emergence of the MTBC, as well as illuminating virulence mechanisms across mycobacterial pathogens as a whole.

Non-tuberculous mycobacteria (NTMs), including *Mycobacterium riyadhense* (MR), are ubiquitous, naturally occurring environmental bacteria commonly found in water and soil[1, 2]. A wide range of animal and environmental sources (aquaria, swimming pools) act as reservoirs for NTMs, and several human disease outbreaks caused by exposure to environmental NTMs have been described[3–6]. With the ability to cause infections in both immunocompromised[7] and immunocompetent[8] individuals, *M. riyadhense* has positioned itself as a clinically important pulmonary pathogen since its discovery in 2009 [2]. The clinical and radiologic characteristics of pulmonary infection caused by *M. riyadhense* are indistinguishable from those caused by *M. tuberculosis*, the most important human pathogen of the MTBC [2, 8].

Similar to *M. tuberculosis*, *M. riyadhense* grows at 37°C and requires 2∼3 weeks[2] to form visible colonies on agar media. However, unlike *M. tuberculosis*, which is a worldwide pathogen transmitted directly from human-to-human with no known environmental reservoirs[9], *M. riyadhense* infections are rare and are transmitted to patients via contact with contaminated water[9] and soil[10], with no evidence of human-to-human transmission yet reported. Infections with *M. riyadhense* have been reported in Asia and Europe in countries including Bahrain, South Korea, France, Italy, and Germany [8,11,12], although most of the recent cases originated in patients from Saudi Arabia[13]. Indeed, the very first case of acute *M. riyadhense* infection was initially misdiagnosed as a case of *M. tuberculosis* infection in a Saudi hospital using commercially available diagnostic tests[14].

It is postulated that *M. tuberculosis* evolved from a free-living environmental ancestor into an obligate pathogen[15, 16]. Our current knowledge indicates that *Mycobacterium canettii* is the most closely related obligate pathogenic species to the MTBC[17, 18]. Infections with *M. canettii* are rare and found solely in people from the Horn of Africa, with no environmental reservoir defined to date[19].

Previous phylogenetic studies have suggested that *Mycobacterium kansasii*[20], *Mycobacterium marinum*[21], *Mycobacterium lacus*[22], *Mycobacterium decipiens*[23], *Mycobacterium shinjukuense*[24] and *M. riyadhense*[25] are closely related to the free-living ancestor of the MTBC based on a single marker gene (e.g., *hsp65*, 16S). In the evolutionary context, a comprehensive understanding of the genomic hallmarks that drive the close phylogenomic relationship of *M. riyadhense* with the MTBC and NTMs is of particular scientific interest.

In this study, we report the characterization of eight clinical isolates of *M. riyadhense*, compare their genomes to members of the MTBC, and provide comprehensive insight into the adaptive changes associated with the evolution of the MTBC from environmental bacteria. We examined the lipid profiles of both rough and smooth variants of *M. riyadhense*, compared them to those of other related mycobacteria, and analyzed the transcriptional response of immunity-related host genes in a murine macrophage infection model with *M. riyadhense*, *M. kansasii*, and *M. bovis* BCG.

Using genome information, we developed a simple PCR-based diagnostic test for the rapid and accurate identification of *M. riyadhense* to minimize the risk of misdiagnosis in a clinical setting.

Our analyses provide a comprehensive description of the hallmarks of *M. riyadhense* that make it one of the closest known environmental relatives of the MTBC, and that can serve to illuminate studies into the evolution and pathogenesis of the MTBC.

## Results

### Clinical manifestation and culture characteristics of *M. riyadhense* isolates

Between April 2011 and March 2017, eight clinical cases of infection with *M. riyadhense* were recorded in male patients aged from 8-82 years (Fig 1). In addition to HIV/AIDS, most of the patients had multiple comorbidities, such as pulmonary and/or systemic hypertension, malignancies, and diabetes mellitus (DM), with the lung being the major site of disease (Fig 1).

**Fig 1.**
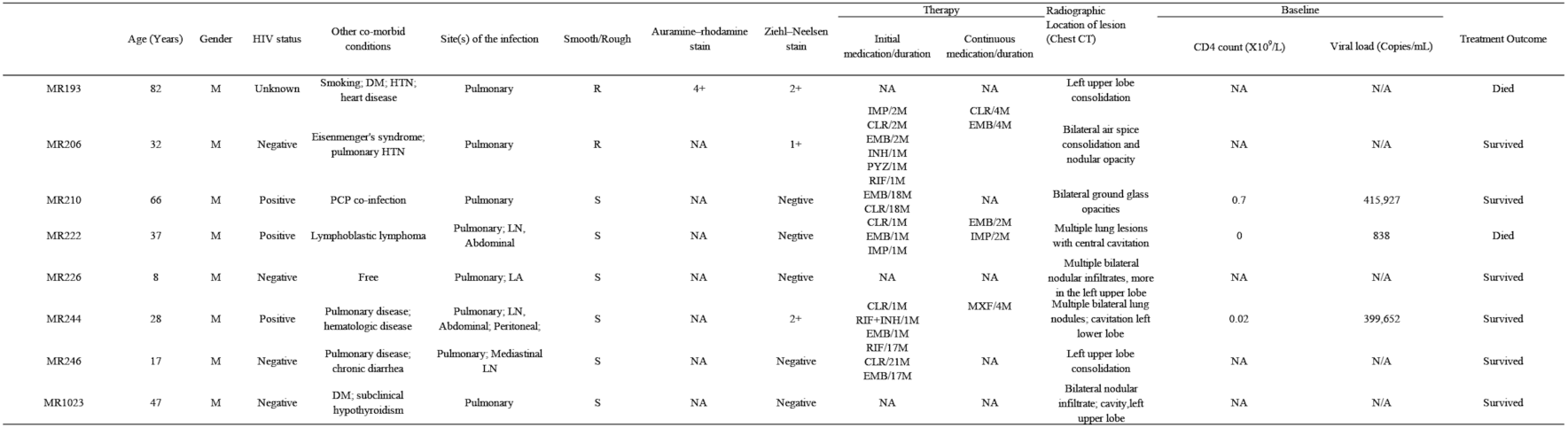
Clinical characteristics of the studies populations.

Bronchoalveolar lavage, endotracheal, sputum, and lymph node biopsy specimens grew *M. riyadhense* colonies when cultured in solid and liquid media using LJ agar and *Mycobacteria* growth indicator tube (MGIT) broth, respectively, with varying times to positivity. Although no consistent findings existed for chest imaging, three out of the eight patients presented with upper lobe consolidation, cavitation, ground-glass opacities, ‘tree-in-bud’ appearance, hilar lymphadenopathy, and pleural effusion (S1 Fig).

The isolates were subjected to susceptibility testing for antibiotics commonly used to treat both typical and atypical infections. Isolates showed 100% susceptibility to rifampin (RIF), rifabutin (RFB), ethambutol (EMB), clarithromycin (CLR), linezolid (LZD), amikacin (AMK), moxifloxacin (MOXI), and trimethoprim-sulfamethoxazole (TMP-SMX). Three out of the eight isolates were resistant to ciprofloxacin (CIP).

### Assembly and annotation of the *M. riyadhense* MR226 genome

The comparison of different assemblies of all sequenced *M. riyadhense* strains is listed in Table S1. We obtained chromosomes of all eight isolates in single contiguous sequences for genome comparison at a high resolution. The *M. riyadhense* MR226 strain contains a 6,243,587 bp chromosome, a linear plasmid (pMRLP01) of 550,247 bp and a circular plasmid (pMR01) of 94,344 bp. The circular nature of the chromosome and the pMR01 plasmid were demonstrated through Gepard[26].

As expected of a free-living opportunistic pathogen, the chromosome of *M. riyadhense* is significantly larger than the chromosomes of the MTBC (Table 1). The number of genes unique in a species showed that the members of the MTBC have a considerably lower percentage of unique genes than *M. riyadhense* and other closely related NTMs (Table 1)

**Table 1.**
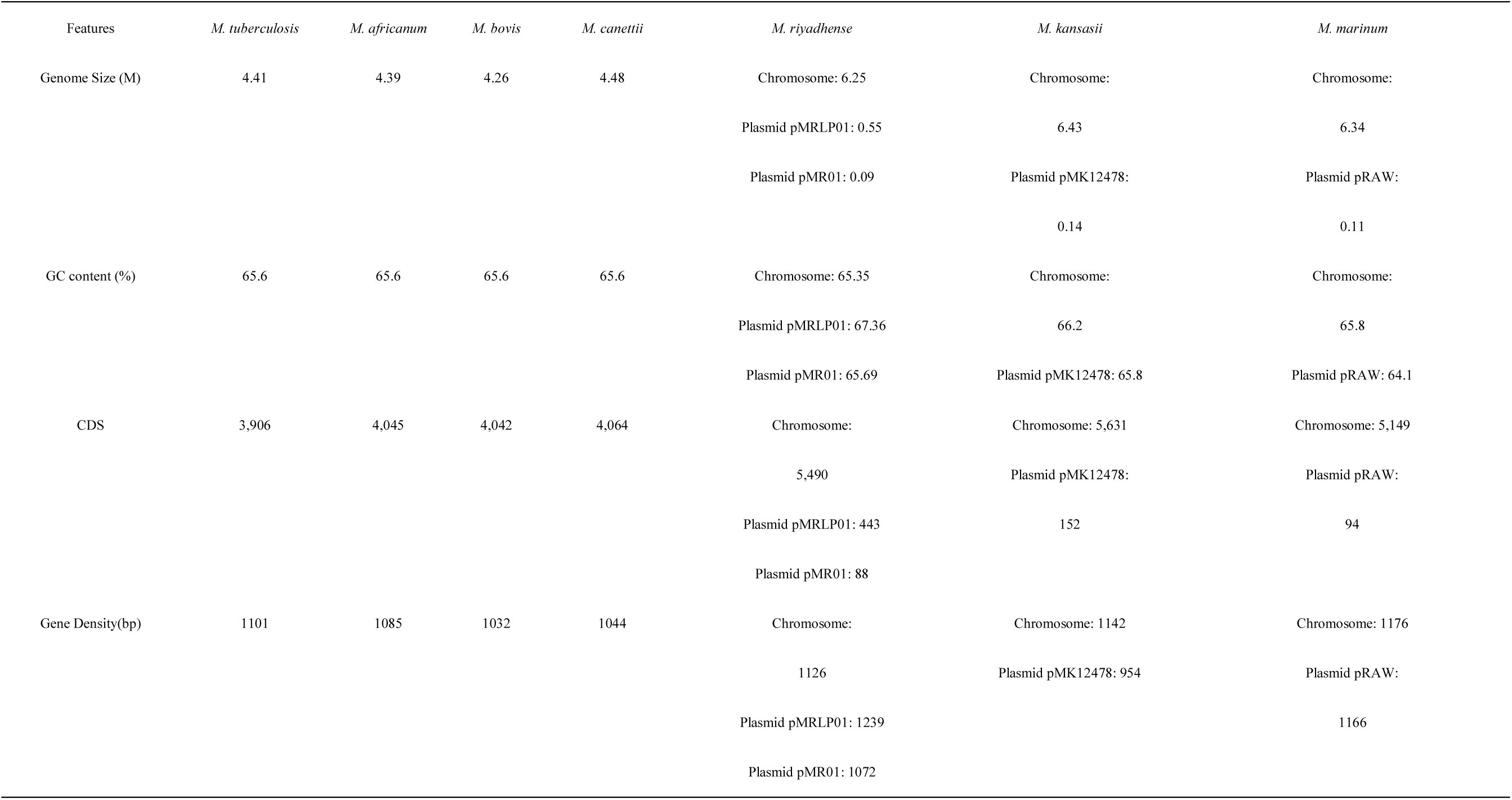
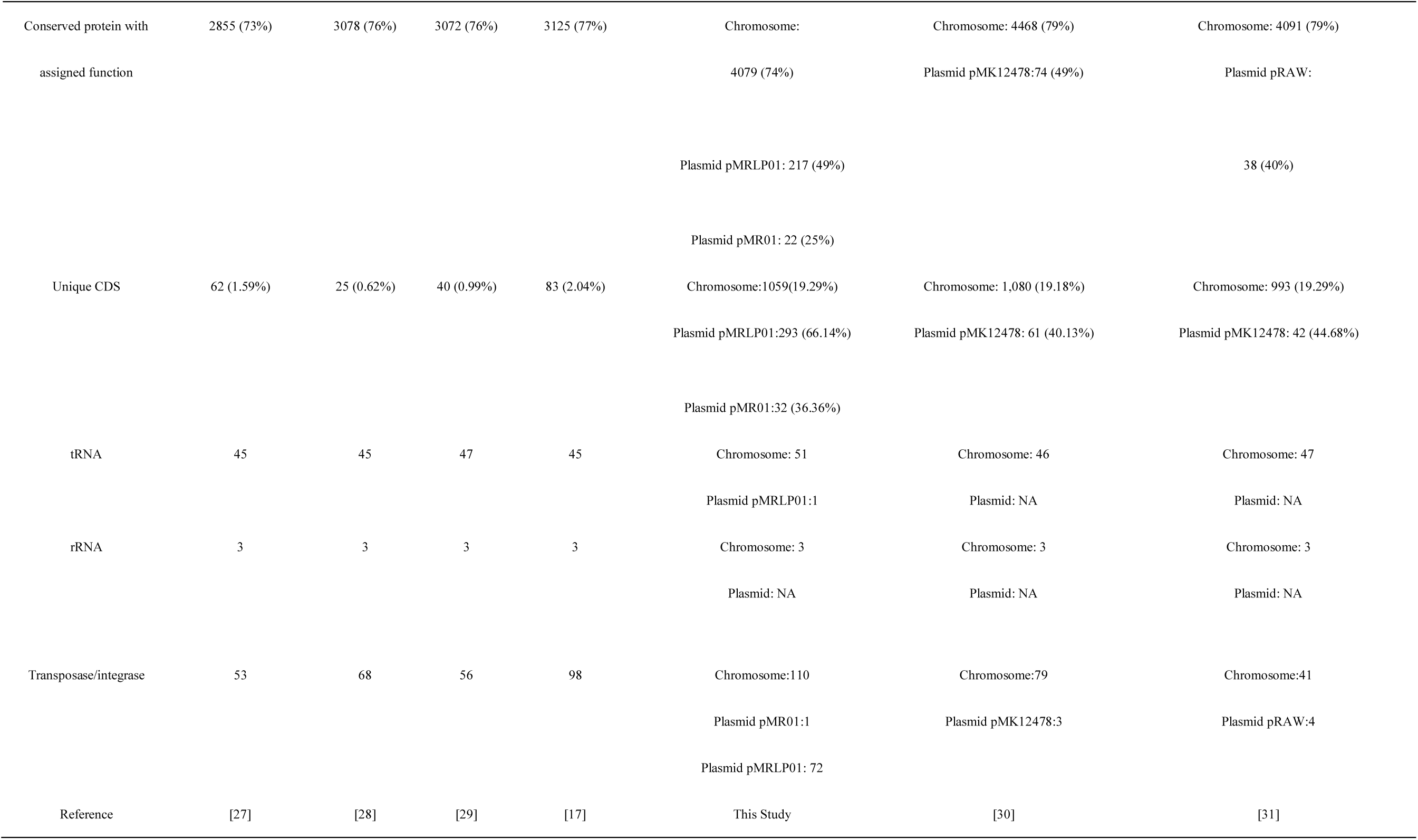
Comparison of general features of genome in Mycobacteria.

| Features | *M. tuberculosis* | *M. africanum* | *M. bovis* | *M. canettii* | *M. riyadhense* | *M. kansasii* | *M. marinum* |
| --- | --- | --- | --- | --- | --- | --- | --- |
| Genome Size (M) | 4.41 | 4.39 | 4.26 | 4.48 | Chromosome: 6.25 | Chromosome: | Chromosome: |
|  |  |  |  |  | Plasmid pMRLP01: 0.55 | 6.43 | 6.34 |
|  |  |  |  |  | Plasmid pMR01: 0.09 | Plasmid pMK12478: | Plasmid pRAW: |
|  |  |  |  |  |  | 0.14 | 0.11 |
| GC content (%) | 65.6 | 65.6 | 65.6 | 65.6 | Chromosome: 65.35 | Chromosome: | Chromosome: |
|  |  |  |  |  | Plasmid pMRLP01: 67.36 | 66.2 | 65.8 |
|  |  |  |  |  | Plasmid pMR01: 65.69 | Plasmid pMK12478: 65.8 | Plasmid pRAW: 64.1 |
| CDS | 3,906 | 4,045 | 4,042 | 4,064 | Chromosome: | Chromosome: 5,631 | Chromosome: 5,149 |
|  |  |  |  |  | 5,490 | Plasmid pMK12478: | Plasmid pRAW: |
|  |  |  |  |  | Plasmid pMRLP01: 443 | 152 | 94 |
|  |  |  |  |  | Plasmid pMR01: 88 |  |  |
| Gene Density(bp) | 1101 | 1085 | 1032 | 1044 | Chromosome: | Chromosome: 1142 | Chromosome: 1176 |
|  |  |  |  |  | 1126 | Plasmid pMK12478: 954 | Plasmid pRAW: |
|  |  |  |  |  | Plasmid pMRLP01: 1239 |  | 1166 |
|  |  |  |  |  | Plasmid pMR01: 1072 |  |  |
| Conserved protein with<br>assigned function | 2855 (73%) | 3078 (76%) | 3072 (76%) | 3125 (77%) | Chromosome:<br>4079 (74%) | Chromosome: 4468 (79%)<br>Plasmid pMK12478:74 (49%) | Chromosome: 4091 (79%)<br>Plasmid pRAW: |
|  |  |  |  |  | Plasmid pMRLP01: 217 (49%) |  | 38 (40%) |
|  |  |  |  |  | Plasmid pMR01: 22 (25%) |  |  |
| Unique CDS | 62 (1.59%) | 25 (0.62%) | 40 (0.99%) | 83 (2.04%) | Chromosome:1059(19.29%)<br>Plasmid pMRLP01:293 (66.14%) | Chromosome: 1,080 (19.18%)<br>Plasmid pMK12478: 61 (40.13%) | Chromosome: 993 (19.29%)<br>Plasmid pMK12478: 42 (44.68%) |
|  |  |  |  |  | Plasmid pMR01:32 (36.36%) |  |  |
| tRNA | 45 | 45 | 47 | 45 | Chromosome: 51<br>Plasmid pMRLP01:1 | Chromosome: 46<br>Plasmid: NA | Chromosome: 47<br>Plasmid: NA |
| rRNA | 3 | 3 | 3 | 3 | Chromosome: 3<br>Plasmid: NA | Chromosome: 3<br>Plasmid: NA | Chromosome: 3<br>Plasmid: NA |
| Transposase/integrase | 53 | 68 | 56 | 98 | Chromosome:110<br>Plasmid pMR01:1<br>Plasmid pMRLP01: 72 | Chromosome:79<br>Plasmid pMK12478:3 | Chromosome:41<br>Plasmid pRAW:4 |
| Reference | [27] | [28] | [29] | [17] | This Study | [30] | [31] |

The comparison of the annotated protein-coding genes from the *M. riyadhense* MR226 strain to the genome assemblies of 152 *Mycobacterium* species (including 77 known mycobacterial plasmids) shows that *M. riyadhense* forms a tight cluster with *M. marinum* and *M. angelicum*, while the MTBC members clusters together with *M. shinjukuense*, *M. lacus* and *M. ulcerans* (S2(A) Fig, highlighted in box a). This clustering is not unexpected as *M. lacus*, *M. shinjukuense* and *M. ulcerans* have smaller genomes and have less predicted proteins as opposed to *M. riyadhense*, *M. marinum* and *M. angelicum*, whose genomes are greater than 6 Mb. A total of 335 genes were identified as unique from the orthologue group comparison, being present in only all sequenced *M. riyadhense* isolates, including MR226. The vast majority of these genes belong to the PE/PPE family, which are thought to be involved in antigen variation and are widely spread across the slow-growing species within the *Mycobacterium* genus.

Linear plasmids were first described in 1989 in maize (which has a linear mitochondrion)[32] and have also been found in *Actinomycetales*, including *Streptomyces* [33], *Rhodococcus*[34] and *Mycobacterium* species, such as *Mycobacterium xenopi*, *Mycobacterium branderi* and *Mycobacterium celatum*. They are often accompanied by a circular plasmid in the same host[35]. The linear plasmid pMRLP01 in *M. riyadhense* contains a pair of partitioning genes (*parA*/*parB*) that are involved in active segregation and thus stabilize the inheritance of the plasmid[36]. The latter are known to contribute to genome evolution by active DNA transfer and exchange[37]. As is often the case for both circular and linear large plasmids, a relatively higher proportion of pMRLP01 genes (51%) have no known function compared to those of the main chromosome (26%). This reinforces the notion that plasmids are an important route by which new genes are introduced into the genome in *Mycobacteria*. Of the 443 predicted protein-coding genes of pMRLP01, 118 have at least one orthologue on the main chromosome. Furthermore, we observed several protein-coding genes in pMRLP01 that have orthologues in the genomes of *Mycobacterium tusciae* JS617, *Mycobacterium aromaticivorans* JCM 16368, *Mycobacterium llatzerense*, *Mycobacterium obuense*, *Mycobacterium novocastrense* and *Mycobacterium holsaticum* (S2(B) Fig). This finding indicates that pMRLP01 can be readily exchanged with other environmental mycobacterial species, with implications for horizontal gene transfer in mycobacteria.

It is well known that plasmids are important “vehicles” for the exchange of genetic material between bacteria or between chromosomes and extra-chromosomal plasmids. In this study, we further identified a circular plasmid termed pMR01 in *M. riyadhense* (Figs S2(C) and S3). When compared with the circular plasmids of other species, such as pRAW in *M. marinum*[38], pMAH135[39] and pMA100[40] of *M. avium*, pMyong1 from *Mycobacterium yongonense*[41], pMK12478[42] from *M. kansasii* and several plasmids from *Mycobacterium chimaera*[43], a high similarity was observed. These plasmids all harbor both a T4S and T7S, which are necessary for conjugation[38, 44], and facilitate the exchange of genetic material between different species of slow-growing mycobacteria[38]. We therefore speculate that pMR01 is a novel conjugative plasmid.

In mycobacteria, five type VII secretion systems have been described, named ESX-1 to ESX-5[45, 46]. An ESX-P5 locus on pMR01, which shows high similarity to the ESX-5 loci on pMK12478, pRAW and pMAH135, is markedly different from the ESX-5 system found on the main *M. riyadhense* chromosome. ESX-5 is involved in the secretion of PE and PPE proteins in *M. tuberculosis* and is involved in modulating the host immune responses to maintain a persistent infection[47]. The potential transmissibility of pMR01 and other pMR01-like plasmids may therefore play a role in the evolution of the ESX systems in mycobacteria.

The progressive alignments (S4(A) Fig) of the assembled chromosomes and plasmids (S4(B) Fig) of each *M. riyadhense* strain show that the chromosomes are relatively conserved; however, the linear plasmids present in all eight sequenced isolates are diverse from both structural and similarity perspectives, while the pRAW-like plasmids are present in only the MR226, MR193 and MR222 strains.

The SNP-based phylogeny of the sequenced *M. riyadhense* isolates based on 43,136 polymorphic sites (S2 Table, S5 Fig) indicates the presence of two different clades of *M. riyadhense* among the clinical isolates sequenced in this study. The nucleotide diversity between the MR222 clade is greater than the diversity between *M. tuberculosis* strains[48], while it is smaller than that seen between *M. canettii* strains[17], and the variation between the MR226 clade is comparable to the SNP variation in *M. tuberculosis* strains.

### Regions of difference (RDs) in *M. riyadhense*

The RDs were originally described as genomic regions present in virulent *M. bovis* and *M. tuberculosis* but absent from the *M. bovis* BCG genome[49]. RD loci were subsequently described across the MTBC[50] and contain functions believed to contribute to pathogenicity[51–53] and evolution of MTBC species[54]. *M. riyadhense* was found to harbor most of the RD loci (RD1, RD3-R11, R13-RD16) that are also intact in *M. tuberculosis,* while 2 of the RDs show unique deletions, RD2^riyadh^ (S6(A) Fig) and RD12^riyadh^ (S6(B) Fig).

RD2 was originally described as deleted in BCG vaccine strains. Subsequently, it was shown that disruption of RD2 in *M. tuberculosis* leads to decreased proliferation *in vivo* and impaired modulation of the innate immune response[52]. The RD2 locus also has a deletion from the *M. riyadhense* genome. It is a larger deletion than the originally described RD2^BCG^ as RD2^riyadh^ contains 29 genes (*rv1971*∼*rv2000*, location 2,216,498∼2,246,766); eight genes within this locus (*rv1978*, *rv1979c*, *rv1980c*, *rv1981c*, *rv1983*, *rv1984*, *rv1987*, *rv1988*) have orthologues elsewhere in the *M. riyadhense* genome (*mr_05764*, *mr_05852*, *mr_02310*, *mr_02993*, *mr_00486*, *mr_02995*, *mr_02325*, *mr_02349*, *mr_01747*), suggesting possible functional redundancy.

The RD12 locus shows deletions across MTBC members, including *M. bovis*, *Mycobacterium caprae* and *M. orygis*[55], but it is present in other MTBC members. *M. canettii* isolates (except group B[56]) also show an independent deletion at the RD12 locus named RD12^can^ (3,479,430∼3,491,866, *rv3111*∼*rv3126*), which is distinct from RD12^bovis^ (3,484,740∼3,487,515, *rv3117*∼*rv3121*). We identified another unique deletion at the RD12 locus in *M. riyadhense*, designated RD12^riyadh^, which encompasses a larger region than RD^can^ and RD12^bovis^, encompassing *rv3108*-*rv3127* (3,477,171∼3,492,150) (S6(B) Fig). It is intriguing that multiple mycobacteria show independent deletion events at the RD2 and RD12 loci, suggesting selective forces play a role in this variation.

### Comparative phylogeny of *M. riyadhense* with other Mycobacteria

The phylogenetic tree shows that the slow-growers and rapid-growers are separated into two different clades and that fast-growers are ancestral compared to slow-growers (Fig 2). The overall topology of our tree is similar to that of previously published phylogenetic trees[22]. In the tree, *M. riyadhense* is located within the same clade as the obligate and opportunistic causal organisms of mycobacterial diseases in humans that include the MTBC, *M. marinum*, *M. kansasii*, *M. leprae* and related host-restricted mycobacteria with reduced genomes and decreased survivability in the environment.

**Fig 2.**
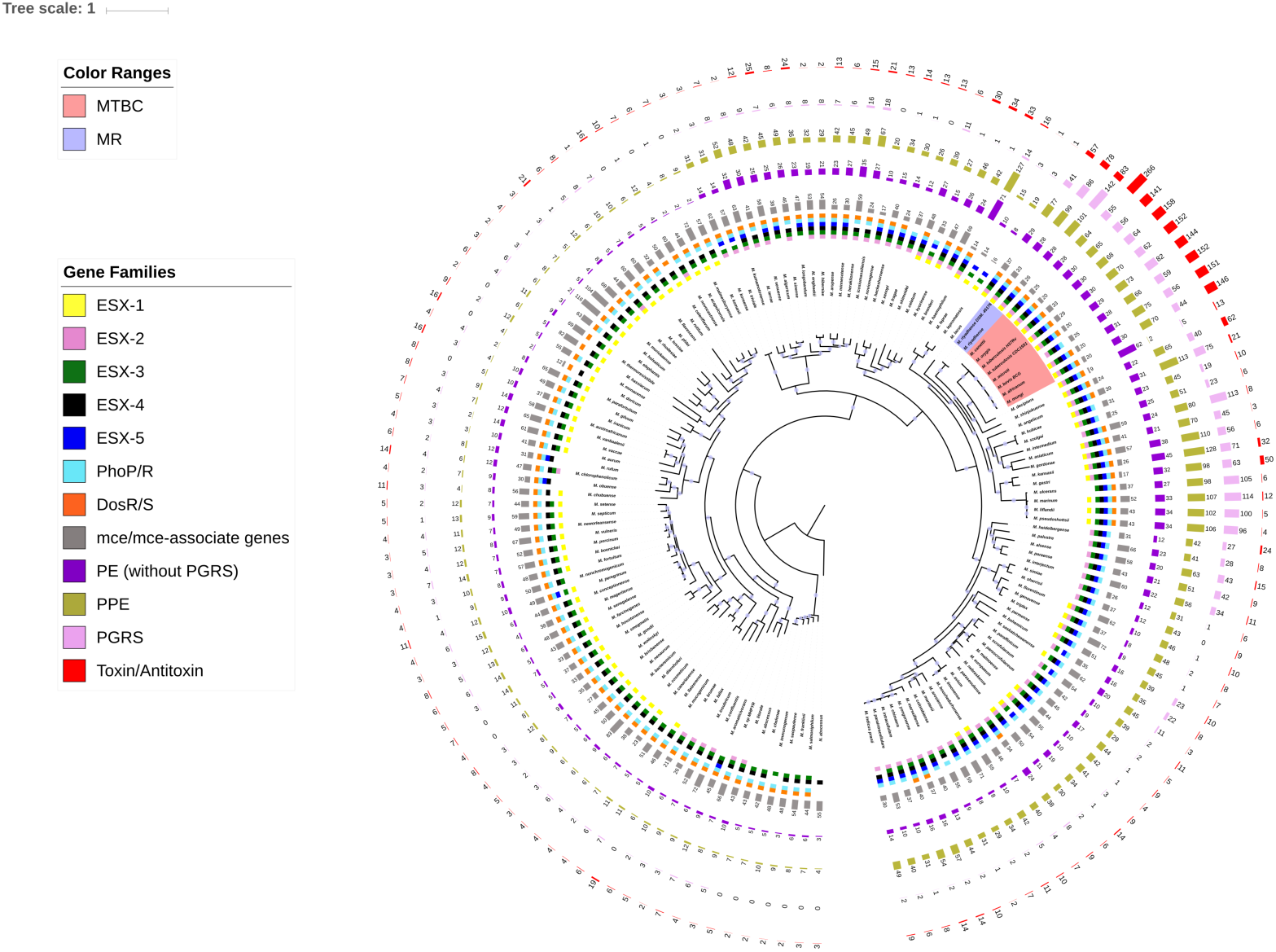
A phylogenetic tree of 149 species of mycobacteria showing close relationship of *M. riyadhense* and MTBC. The phylogeny is constructed using 149 available genomes by concatenating and aligning amino acid positions across 400 shared proteins automatically identified in the chosen genomes by PhyloPhlAn2. The description of the gene families is listed on the left side. For details please refer to the Materials and Methods. Circles on each branch indicate the bootstrap values above 95%.

The PE/PPE family *mce* and *mce*-associated genes are known to be important for host adaptation[57] and pathogenicity[58]. The PE/PPE family genes are enriched in the MTBC members but also in *M. riyadhense* MR226 (278) and other pathogenic species, such as *M. kansasii* (228) and *Mycobacterium ulcerans* (200). The number of *mce-* or *mce*-associated genes has not significantly changed across mycobacterial genomes, indicating that this group of genes is under an evolutionary constraint and plays functional roles bridging both environmental and obligate pathogen lifestyles. Our results agree with previous findings that during their evolution, the ESX systems were derived from the ancestor ESX-4, as shown in Fig 2 at the root node, and then ESX-3, ESX-1, ESX-5 and ESX-2 evolved by horizontal transfer[59].

A comparative phylogenetic map based on 906 conserved proteins (S7 Fig) reveals this downsizing of the genome and the dynamic changes in genome components. Certain functional categories of genes are relatively enriched during the evolution of MTBC, including protein metabolism, regulation and cell signalling, cell division and cell cycle.

*M. riyadhense* shares a larger number of orthologues (3,122) with *M. tuberculosis* than with *M. kansasii* (2,978), *M. marinum* (2,962) and *M. szulgai* (2,724) among the environmental mycobacteria that are closely related to the MTBC (Fig 3(A)). A total of 134 orthologues are exclusively shared between *M. riyadhense* and *M. tuberculosis*, while the number of orthologues exclusively shared between *M. tuberculosis* and *M. kansasii* (30), *M. marinum* (48) and *M. szulgai* (18) is less (Fig 3(A)). It is notable that genes from the phage-derived regions of RD3 and RD11 are shared exclusively between *M. riyadhense* and *M. tuberculosis*.

**Fig 3.**
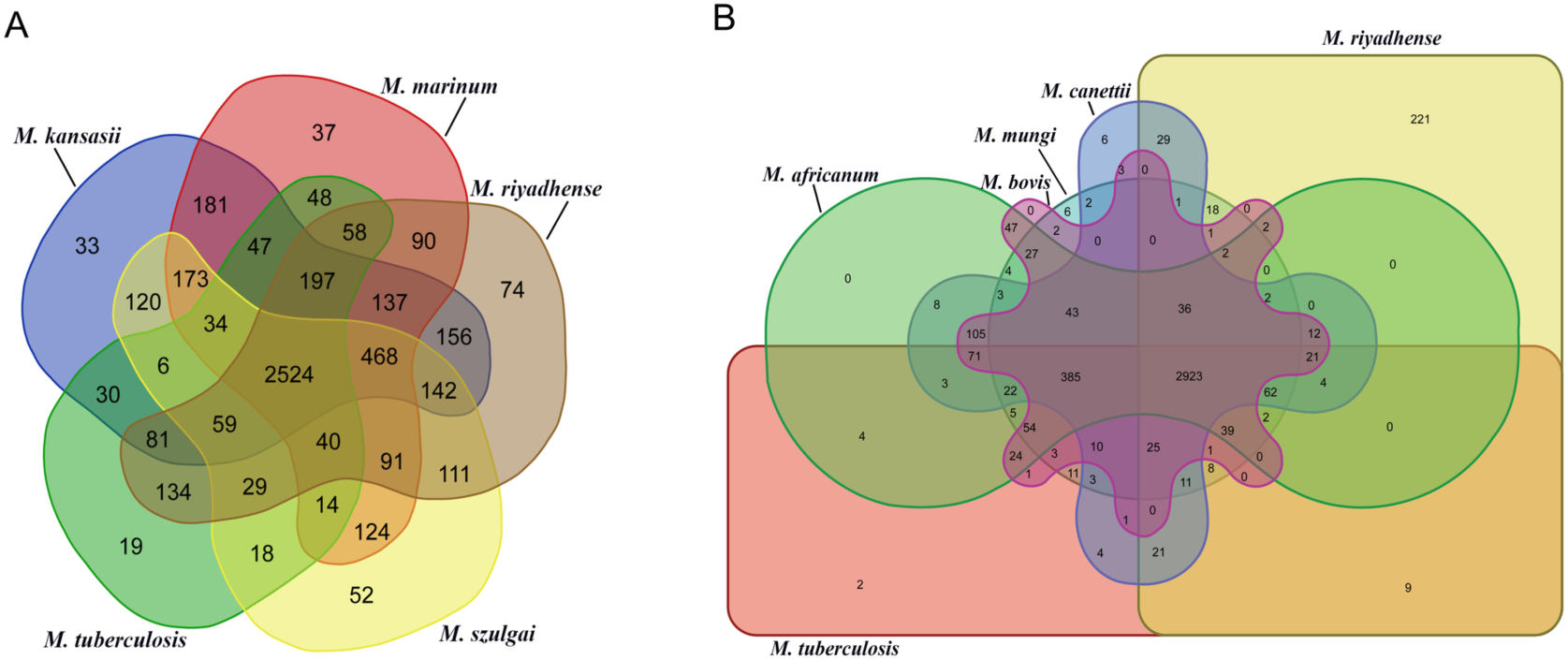
Comparison of the shared and unique gene orthologs amongst selected mycobacteria. Venn diagrams represent the overlap of gene orthologs between the (A) *M. riyadhense*, *M. tuberculosis* H37Rv, *M. marinum* M, *M. kansasii* 12478 and *M. szulgai* and (B) *M. riyadhense* MR226 and five species within the MTBC.

The comparative analysis of the orthologue groups of *M. riyadhense* and the MTBC is informative. Firstly, 385 orthologue groups present across all MTBC species are absent from *M. riyadhense*. Secondly, 221 orthologues uniquely present in *M. riyadhense* are not found amongst the MTBC (Fig 3(B)). This latter group of *M. riyadhense* unique orthologues likely depict the constraints imposed by the free-living biology of *M. riyadhense* which necessitates maintaining a broad functional repertoire to secure environmental survival. In contrast, the obligate MTBC species have lost genes involved in environmental survival but gained a large number of genes required for survival in the *in vivo* environment.

A hallmark of *M. tuberculosis* infection is the ability to survive long-term in host granulomas and develop a latent stage of infection. The molecular mechanisms and cellular components that are involved in the persistence of *M. tuberculosis* are still poorly understood, but several T/A systems have been implicated in the pathogenicity of *M. tuberculosis*[60]. T/A systems were first found on plasmids or plasmid-derived chromosomal loci where they promoted plasmid maintenance in bacterial populations[61], but when compared to other mycobacteria, the MTBC are remarkable for the extensive expansion of T/A systems. We thus compared the 79 pairs of T/A systems (belonging to the HigAB, MazEF, ParDE, RelEF, VapBC and UCAT families) in *M. tuberculosis* with the T/A pairs found in other members of the MTBC and NTMs. Based on the presence of 49 out of the 79 T/A orthologue pairs (Fig 4), *M. riyadhense* appears more closely related to the MTBC than to other mycobacteria, including *M. lacus, M. shinjukuense* and *M. decipiens*. It can be hypothesized that the shared component of T/A pairs play a functional role in the pathogenicity or persistence of *M. riyadhense* infection in a way similar to that described for the importance of T/A pairs in *M. tuberculosis in vivo* biology.

**Fig 4.**
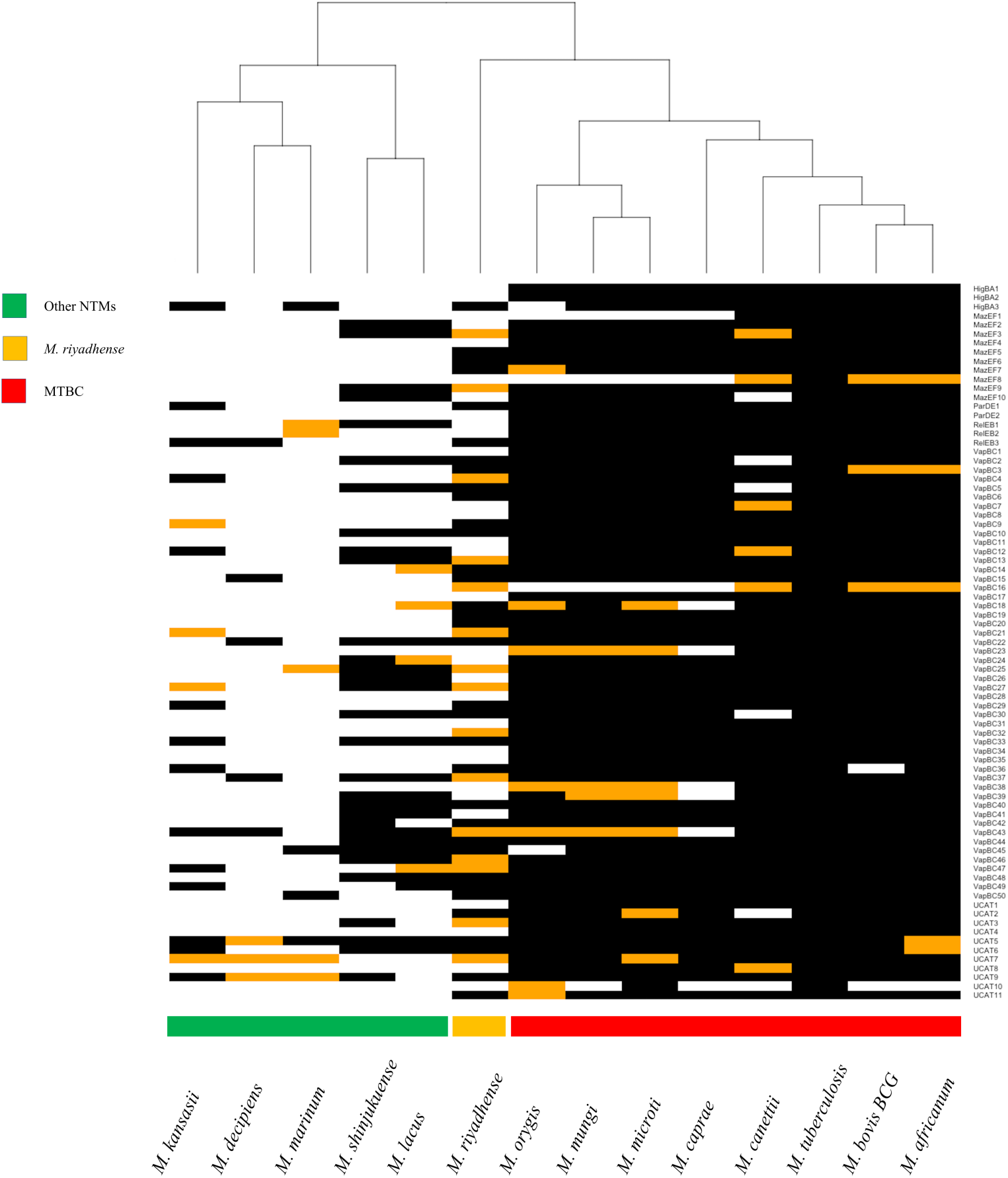
A hierarchical clustering of the presence (black) and absence (white) of *M. tuberculosis* H37Rv toxin/antitoxin orthologs in *M. riyadhense*, *M. marinum*, *M. kansasii*, *M. shinjukuense*, *M. lacus*, *M. decipiens* and MTBC species. The orange blocks denote the presence of either the toxin or antitoxin gene ortholog in a given pair of the T/A system. The black and white blocks represent presence and absence respectively. The name of the T/A system are shown for each row on the right.

### *M. riyadhense* strains produce a distinct pattern of LOSs

The lipopolysaccharides (LOSs) are an important class of glycolipids that have previously been linked to diverse mycobacterial phenotypes, such as colony morphology, secretion of PE/PPE family proteins, and the pathogenicity of *M. marinum*[62]. It is noteworthy that we observed both smooth (MR210, MR22, MR226, MR244, MR246, and MR1023) (Fig 5(A)) and rough (MR193 and MR206) (Fig 5(B)) morphologies in *M. riyadhense* strains. We therefore sought to first examine whether the genetic machinery for the production of LOS is present in the *M. riyadhense* genome, and then to follow up on the genome-level predictions with lipid analyses of rough and smooth variants using Thin-layer Chromotagraphy (TLC).

**Fig 5.**
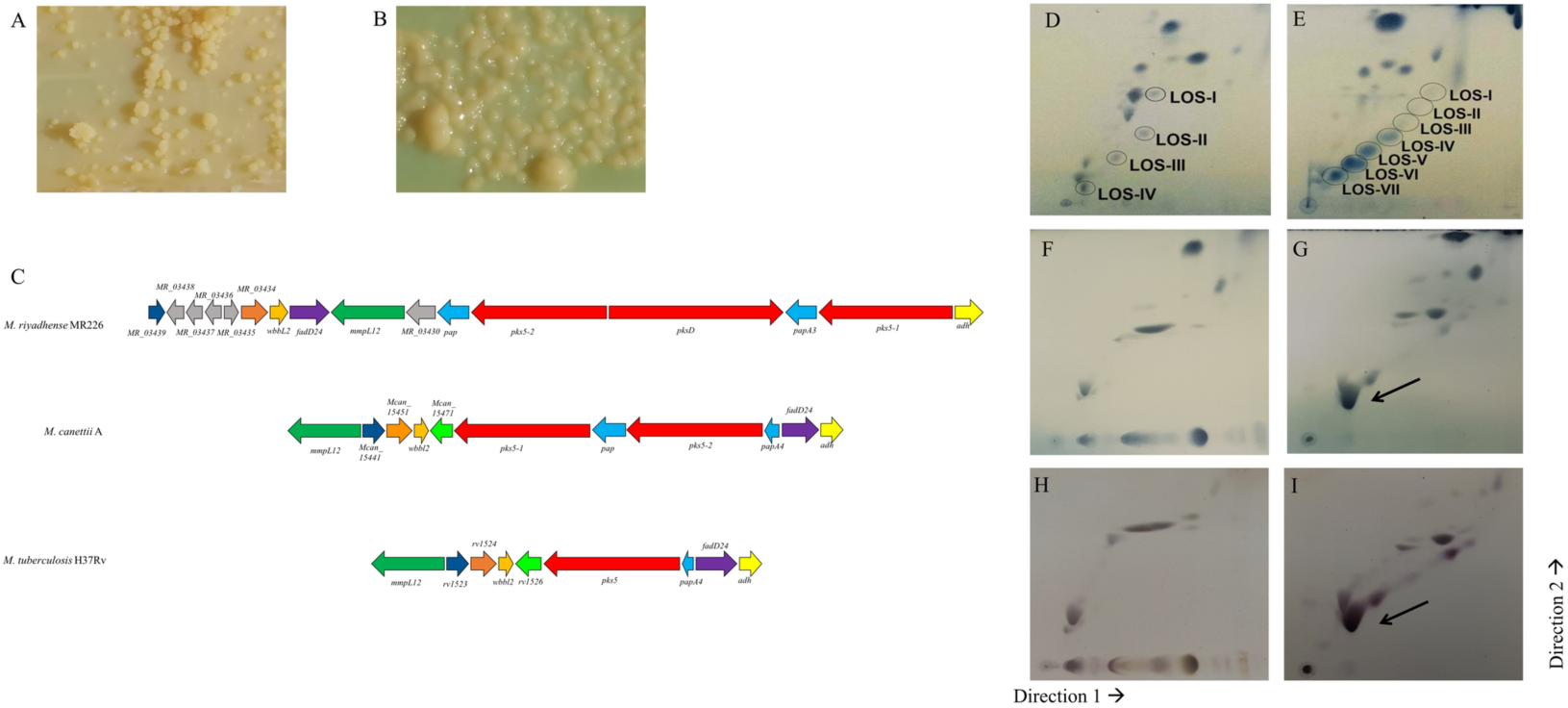
LOS systems in *M. riyadhense* and other related mycobacteria and 2D-TLC analysis of polar lipids extracted from selected *M. riyadhense* strains. (A) Rough-dry colony morphology (MR193) and (B) smooth morphology (MR226) of *M. riyadhense*. (C) Genetic locus map of the LOS biosynthesis gene cluster from *M. riyadhense*, *M. canettii* A and *M. tuberculosis* H37Rv (drawn to scale). The arrows show the direction of transcription and the genes are colored according to the orthologous relationships. Polar lipids from two known LOS producers, *M. marinum* (D) and *M. kansasii* (E), are included to illustrate the migration pattern of LOS species in System E. (F) 2D-TLC analysis of polar lipids extracted from select *M. riyadhense* rough strain or smooth (G) strain. A separate staining with alpha-napthol also confirmed that this was a glycolipid species from the same (H) rough and (I) smooth strain. (D), (E), (F) and (G) were charred after staining with MPA, while (H) and (I) were charred after staining with alpha napthol. LOS III from *M. riyadhense* is indicated by a solid arrow.

We observed that the *wecE* and *galE6* LOS genes are absent from the *M. riyadhense* genome (Table 2, S3 Table). These genes are linked with the removal of LOS II* and the production of LOS IV[62]. Thus, their absence is likely to cause an accumulation of LOS II* and the lack of fully formed LOS IV, which have previously been shown to increase the pathogenicity of *M. marinum*[62]. Furthermore, both the *pks5* and *pap* genes in the LOS locus are intact in *M. riyadhense*, as is the case in *M. canettii*, but not in *M. tuberculosis,* where the former is truncated and the latter deleted[63]. This finding indicates that *pks5* recombination and *pap* deletion occurred in a common ancestor of the MTBC after its differentiation from both *M. riyadhense* and *M. canettii.* Remarkably, the *M. riyadhense* LOS gene locus layout is dissimilar to that in *M. canettii*, *M. tuberculosis*, *M. kansasii* and *M. marinum*: indeed, exclusive rearrangements of this locus in *M. riyadhense* were observed (Fig 5(C)).

**Table 2.** Presence and absence of the *M. marinum* orthologs related to LOS synthesis in *M. riyadhense* and other species

|  | *M. smegmatis* | *M. kansasii* | *M. canettii* | *M. riadhense* | *M. tuberculosis* |
| --- | --- | --- | --- | --- | --- |
| MMAR_1008 | P | P | P | P | P |
| MMAR_2307 | A | P | A | P | A |
| MMAR_2313 | A | P | P | P | P |
| MMAR_2319 | A | A | A | A | A |
| MMAR_2320 (*wecE*) | A | P | P | A | A |
| MMAR_2327 | A | P | P | P | P |
| MMAR_2332 | A | A | A | A | A |
| MMAR_2336 (*galE*) | A | P | A | A | A |
| MMAR_2340 (*pks*) | P | P | P | P | T |
| MMAR_2341 | P | P | P | P | P |
| MMAR_2343 (*pap*) | P | P | P | P | A |
| MMAR_2353 | P | P | A | P | A |
| MMAR_5170 (*whiB4*) | P | P | P | P | P |

P: Present; A: Absent; T: Truncated

To correlate rough vs. smooth colony morphology with LOS production, we extracted polar lipids from the strains and analyzed them by 2D-TLC using solvent system E[64], which is designed to separate phospholipids and LOSs. Charring of the TLC plates with alpha-napthol revealed glycolipids, including the accumulation of a species that migrated at a position similar to that of LOS III. This lipid was seen in only smooth strains; species with migration patterns similar to LOS I and LOS II were observed, while LOS IV was absent. This result was not unexpected because all *M. riyadhense* strains lack a functional *wecA* gene, which is required for the extension of LOS II to LOS IV (Fig 5). Additionally, the relative levels of the predominant LOS species in *M. riyadhense* seem to be quite high when compared to those seen in other LOS-producing mycobacteria (Figs 5(D)(E)). Conversely, the rough strains did not produce any glycolipids that migrated in the positions corresponding to LOSs (Figs 5(F)(H)).

### *PE-PGRS33* locus and type VII secretion system of *M. riyadhense*

The *pe-pgrs33* (*rv1818c*) locus encodes the exported protein PE_PGRS33 and plays an important role in the pathogenesis of *M. tuberculosis*[65]. A previous study[66] showed that *pe-pgrs33* is present in all MTBC members but not in *M. canettii*, which implies a specific *pe-pgrs33* insertion event in the ancestor of MTBC strains. Genome comparison of *M. riyadhense* with *M. tuberculosis*, *M. kansasii*, *M. marinum* and *M. canettii* provides additional evidence that *M. riyadhense* is the missing link of the *pe-pgrs33* deletion/insertion event. Our phylogeny strongly suggests that the deletion of *pe-pgrs33* from *M. kansasii* and *M. marinum* occurred before the divergence of environmental mycobacteria and the smooth tubercle bacilli (STB)/MTBC clade (S8 Fig).

All 5 ESX systems (ESX1-ESX5) were found in the *M. riyadhense* genome (S9 Fig) but with minor modifications. Hence, the *eccA* and *eccB* genes are absent from the ESX-2 system, while the *espACD* operon, which is essential for secretion of virulence factors via ESX1, is also missing in *M. riyadhense*. The overall gene arrangement of the ESX1-ESX5 loci is similar in both *M. riyadhense* and *M. tuberculosis*[45]. This conserved synteny reinforces the previous results that phylogenetically *M. riyadhense* may represent an ancestral state to MTBC. As noted before, the pMR01 plasmid also contains an extra ESX-P5 locus, which could indicate a role for this plasmid in mediating pathogenicity (S3 Fig).

### Transcriptional response of murine macrophage cells upon *M. riyadhense* infection

Our genomic analysis of *M. riyadhense* revealed a range of genes and potential systems that could play a role in host-pathogen interactions. We therefore sought to assess the initial interaction of *M. riyadhense* with macrophages, using the RAW264.7 cell line of murine origin as our experimental model. As comparator strains in our analysis, we performed parallel infections with *M. kansasii*, an opportunistic pathogen that also contains an orthologous ESX-1 system (Fig 2), and *M. bovis* BCG, the live TB vaccine that is attenuated through deletion of the ESX-1 system. These comparisons allowed us to explore whether *M. riyadhense*-triggered innate immune responses were more similar to those triggered by *M. kansasii* or BCG or were intermediate between the two.

To analyze the innate immune responses of the macrophages in the intracellular presence or absence of these mycobacterial isolates, the transcriptional response was analyzed using a 754 probe NanoString Murine Myeloid Innate Immunity panel V2[67] at 3, 24 and 48 hours post infection. These analyses revealed an expected commonality in the responses to infection with all mycobacteria, such as induction of proinflammatory genes through TLR signalling (e.g., upregulation of *IL1B*, *TNFA*, *CCL4*, *PTGS2* and *CXCL2*, albeit to different absolute levels, (Figs 6, S10(A) and S10(B)). Distinct responses triggered by BCG infection compared to *M. riyadhense* and *M. kansasii* included, for example, upregulation of *MARCO* by BCG infection (S10(B) Fig); *MARCO* is involved in pathogen uptake via trehalose dimycolate, a lipid that is known to show variation in structure between the MTBC and *M. kansasii*[68, 69]. *CCL24* and *CXCL14*, which are involved in the attraction of immune cells to the site of infection[70], were upregulated to higher levels with *M. riyadhense* and *M. kansasii* than with BCG at 24 and 48 hours.

Overall, our analysis of the macrophage transcriptional profiles showed that the response to infection with *M. riyadhense* and *M. kansasii* triggered more similar macrophage transcriptional responses in comparison to the response produced by BCG infection (S10 Fig).

### Developing a rapid PCR-based diagnostic marker for *M. riyadhense*

Due to the issues previously encountered in diagnosing *M. riyadhense* infections[2, 8], correct and prompt identification of cases upon presentation at healthcare units is of paramount importance. We therefore sought to translate our knowledge on the genome sequences into a PCR diagnostic test that could be used in a clinical microbiology setting to distinguish *M. riyadhense* from other mycobacteria, including the members of the MTBC.

By identifying unique K-mers ranging in size from 11 bp to 4,209 bp (S11 Fig) in the assembled genome compared to the genomes of 152 other mycobacterial species, four primer sets were developed targeting the *mr_00036*, *mr_00263*, *mr_00606* and *mr_01005* genes. The MRDP primer pair MRDP-F/MRDP-R amplified a single product from each of the eight isolates of *M. riyadhense* (Fig 7) but not from other mycobacterial species, including *M. tuberculosis*, *M. bovis*, *M. kansasii*, *M. marinum*, *M. szulgai*, *M. avium* and *Mycobacterium angelicum*. This result shows that the MRDP-F/MRDP-R primers are highly specific to *M. riyadhense* and form the basis for a simple diagnostic PCR that can inform appropriate treatment protocols.

**Fig 6.**
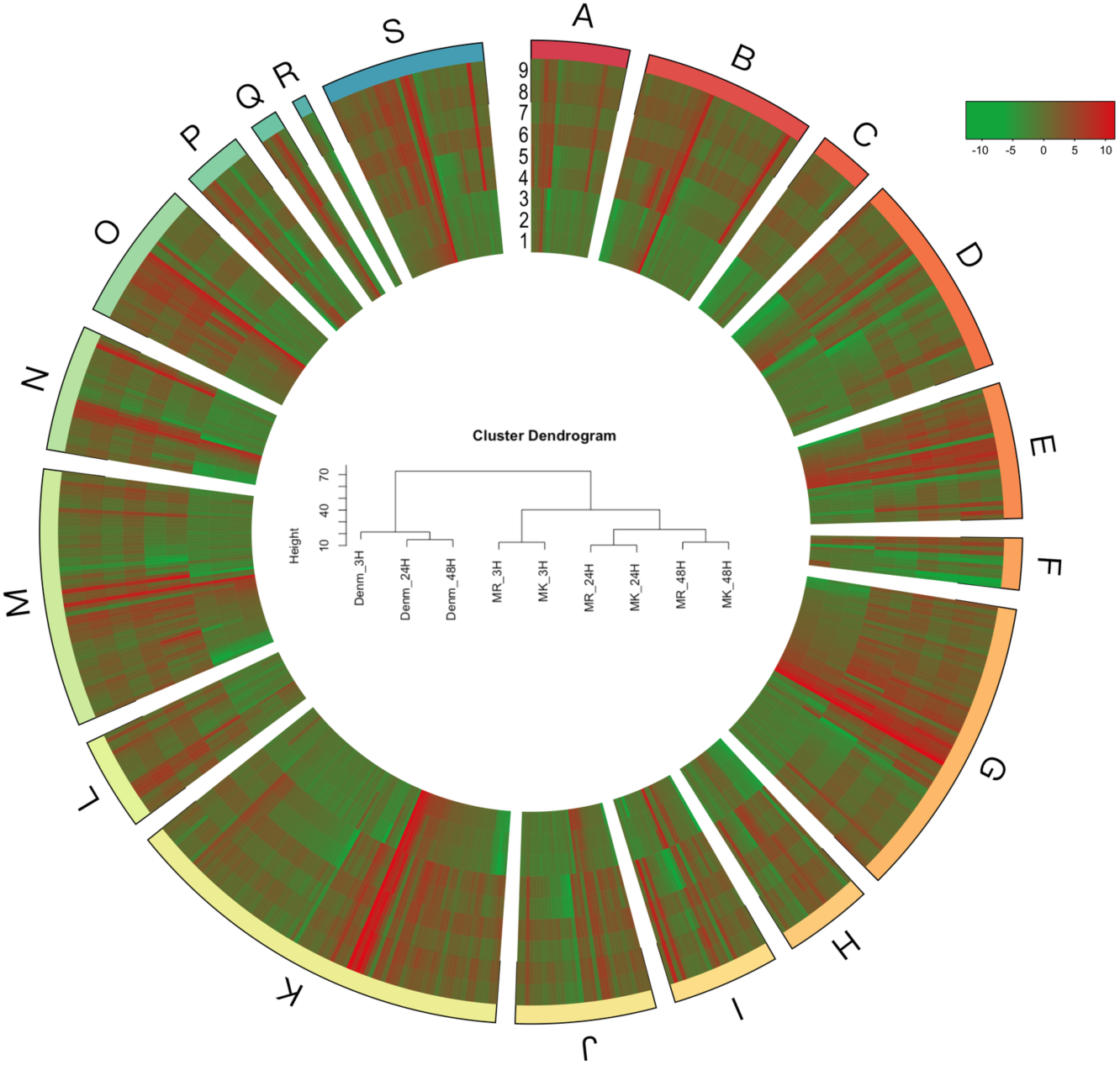
A circular heatmap demonstrating the host gene expression patterns from the RAW264.7 infections with *M. riyadhense* (MR), *M. kansasii* (MK) and *M. bovis* BCG Denmark (Denm) over 3, 24 and 48 hours as determined by the Myeloid Innate Immunity Gene Expression Panel (NanoString Technologies, USA). Data represent Log2FC expression values compared to the uninfected control from each time point. The gene sets of each clusters are: A: Angiogenesis; B: Antigen Presentation; C: Cell Cycle and Apoptosis; D: Cell Migration and Adhesion; E: Chemokine signaling; F: Complement Activation; G: Cytokine Signaling; H: Differentiation and Maintenance of Myeloid Cells; I: ECM remodeling; J: Fc Receptor Signaling; K: Growth Factor Signaling; L: Interferon Signaling; M: Lymphocyte activation; N: Metabolism; O: Pathogen Response; P: T-cell Activation and Checkpoint Signaling; Q: TH1 Activation; R: TH2 Activation; S: TLR signaling. The circles 1-9 represent RAW264.7 transcriptome response upon infections of 1: Denm-3h; 2: Denm-24h; 3: Denm-48h; 4: MR-3h; 5: MK-3h; 6: MR-24h; 7: MK-24h; 8: MR-48h; 9: MK-48h. The cluster dendrogram in the middle show the hierarchical clustering of the individual gene expression datasets.

**Fig 7.**
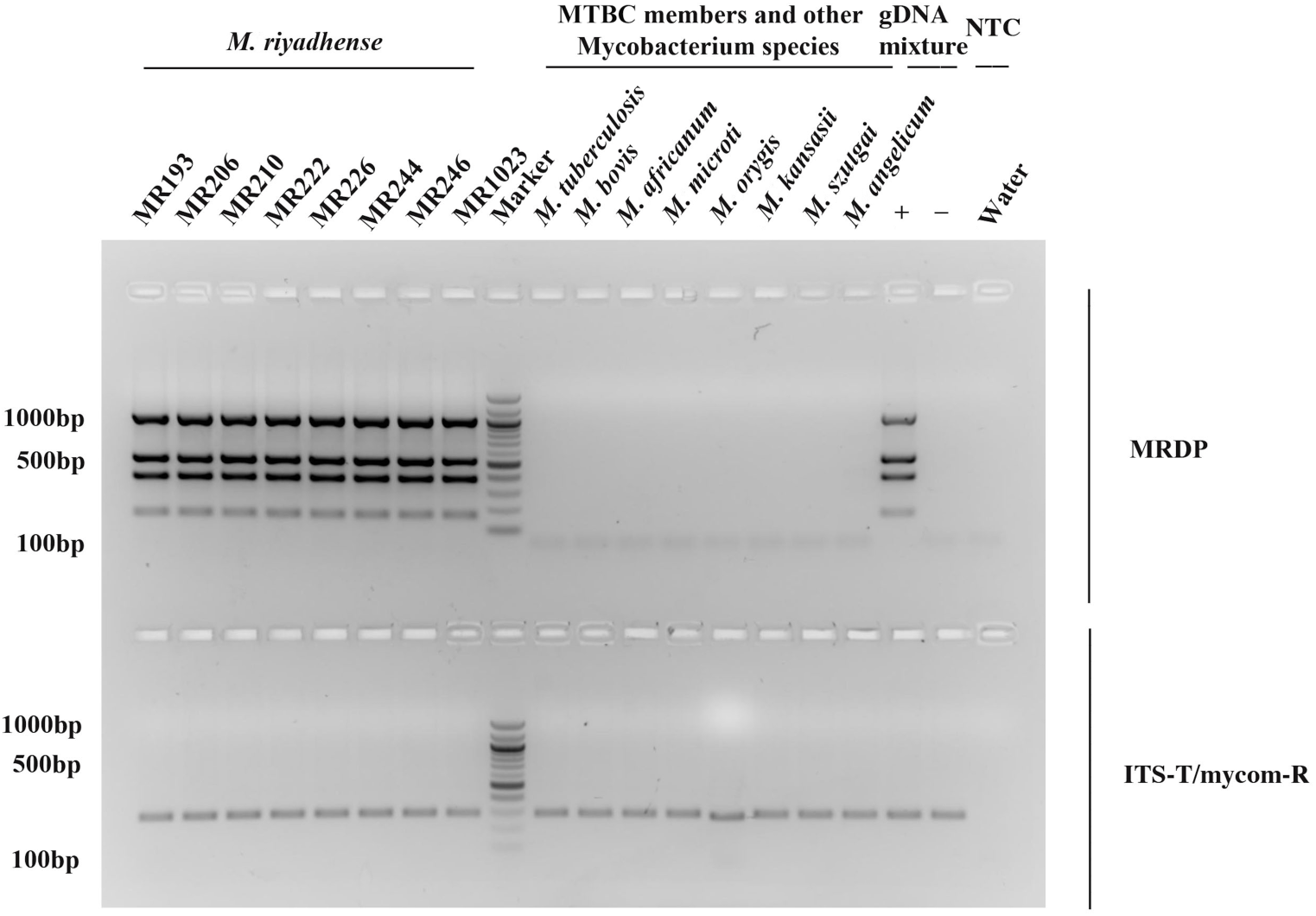
Development of a rapid PCR-based diagnostic test for detection of *M. riyadhense*. Agarose gel (2%) electrophoresis patterns of the PCR products are shown as part of the diagnostic test for *M. riyadhense*. Lane M: DNA Marker, Lane 1-8: Varies *M. riyadhense* strains (From left to right: MR193, MR206, MR210, MR222, MR226, MR244, MR246, MR1023), Lane 9-16: Varies *Mycobacterium* species (From left to right: *M. tuberculosis*, *M. bovis*, *M. africanum*, *M. microti*, *M. oryis*, *M. kansasii*, *M. szulgai* and *M. angelicum*) templates, Template cocktail, mycobacterium species (*M. tuberculosis*, *M. bovis*, *M. kansasii*, *M. marinum*, *M. szulgai*, *M. avium* and *M. angelicum*) with (Lane 17, +) and without (Lane 18,-) *M. riyadhense* MR226 gDNA template. Lane 19: Non-template control (NTC). Upper Panel: MRDP (*M. riyadhense* diagnostic marker) set. Lower Panel B: *Mycobacterium* genus specific primer ITS-T and mycom-R amplified with mycobacterial gDNA.

## Discussion

*M. riyadhense* has become a clinically relevant NTM species globally[8, 71]. Contrary to prior publications on *M. riyadhense* that have been based primarily on clinical case reports, here we present the largest and most comprehensive genomic study undertaken to date on clinical *M. riyadhense* isolates. The eight new *M. riyadhense* strains sequenced in this study originated from pulmonary infections with some having additional extra-pulmonary involvement, which fulfilled the American Thoracic Society/ Infectious Diseases Society of America (ATS/IDSA) criteria for NTM infection[72].

Our comparative analysis of *M. riyadhense* genomes with the MTBC and a large collection of NTMs provide unequivocal evidence that *M. riyadhense* is one of the closest known environmental mycobacteria species to the MTBC and forms a phylotype with *M. lacus*, *M. decipiens* and *M. shinjukuense*. Indeed, while our manuscript was in preparation, independent work [73] also showed the close phylogenetic relationship of *M. riyadhense* to the MTBC, suggesting that it forms part of an MTB-associated phylotype. Our analyses of multiple *M. riyadhense* isolates complements and extends the findings of Sapriel and Brosch by revealing that expansion of T/A pairs, modification of secretion systems, alterations in cell wall lipids, and plasmid-mediated horizontal acquisition of new functionality all played key roles in the evolution of the MTBC. Our work hence adds new insight into the evolution of the MTBC from free-living environmental bacteria to obligate pathogens.

Our study shows that *M. riyadhense* shares a larger number of orthologues with *M. tuberculosis* than *M. kansasii* and *M. marinum*, notably in the T/A gene family (Figs 3 and 4). Forty-nine pairs of T/A gene orthologues were found in *M. riyadhense*, far greater than the number of orthologues observed in any of the other NTMs. The expansion of T/A genes among the MTBC offers additional evidence that suggested original acquisition of T/A modules into mycobacteria through lateral gene transfer played a key role in the development of pathogenicity[21, 74].

*M. riyadhense* strains appeared as both smooth and rough colony forms when grown on solid LJ media. The observation of smooth and rough colony variants is seen in other mycobacterial pathogens where it is linked to presence or absence of LOS; for example, the presence or absence of LOS from *M. canettii* causes a transition from smooth to rough colony variants, respectively, with rough variants showing increased virulence; MTBC strains lack LOS, and it has been suggested that the removal of LOS was a key event in the evolution of the MTBC species towards their current obligate pathogen status [63]. In *M. riyadhense* smooth colony variants we observed the presence of LOS I and LOS II but the absence of LOS IV. These biochemical observations agree with the genomic prediction that *M. riyadhense* strains lack a functional *wecA* gene, which is required for the extension of LOS II to LOS IV. Overall, our results show that *M. riyadhense* exhibits a LOS production phenotype distinct from that of other LOS-producing mycobacteria.

The ESX Type VII secretion systems are key elements of mycobacterial virulence. All the 5 ESX type VII secretion systems present in *M. tuberculosis*, and known to be involved in virulence and pathogenicity, were found in *M. riyadhense* with very similar gene arrangement. An additional ESX-P5 system was also found on the circular plasmid pMR01. pMR01-like plasmids have been shown in many different NTMs, including *M. kansasii*, *M. marinum*, *M. chimaera* and *M. avium*, indicating that these groups of plasmids containing both T4S and T7S are conjugative plasmids. The extensive presence of these plasmids may also explain the origin of the type VII secretion system [38]. ESAT-6 and CFP-10, which are secreted through ESX-1, are critical to phagosome perturbation and manipulation of host macrophages induced by *M. tuberculosis*[75]. Linked to this, our transcriptome analysis revealed that *M. riyadhense* and *M. kansasii* trigger similar overall macrophage responses after infection when compared to *M. bovis* BCG (as a representative of the MTBC). These data reinforce the phylogenetic relationship of *M. riyadhense* with the MTBC species and its transitional status between an opportunistic and an obligate pathogen.

The clinical presentation of our cases was by and large indistinguishable from disease caused by *M. tuberculosis*, as reported earlier[7, 8], but with a negative *M. tuberculosis* PCR. Due to the relatively recent emergence of *M. riyadhense* as an important clinical pathogen coupled with its misdiagnosis as *M. tuberculosis* by commercially available kits, we developed an accurate set of diagnostic markers based on the genomic datasets generated in this study. The primer sets accurately detect *M. riyadhense* in a mixed cocktail of closely related mycobacteria, and can hence serve as part of an accurate and fast diagnostic protocol in clinical settings thus reducing the need for strict isolation, laborious contact tracing, and inappropriate use of TB antimicrobials. Going forward, these primers could be used in a global epidemiological survey of cases of *M. riyadhense* infections in humans, animals and the environment, providing a more complete picture of the epidemiology of *M. riyadhense* following a broad One Health approach. It would, for example, be of clinical interest to see if *M. riyadhense* infections occur in Africa and South America, for which no reports are available, or if *M. riyadhense* is uniquely endemic in the Arabian Peninsula. Indeed, a limitation of this and previous similar studies is that all isolates available for genomic studies are clinical. *M. riyadhense* is an environmental pathogen, and hence it may be found in animals sharing the same niche with humans that were infected from the environment. As the environmental reservoir of *M. riyadhense* remains unknown, we believe that systematic screening of relevant environmental samples with the MRDP established in this study may help to establish the natural habitat of this bacterium and hence allow improved infection control.

In conclusion, our study provides unprecedented insights into the ancestry and adaptive evolution of the MTBC relative to extant NTM species and places *M. riyadhense* as one of the closest relatives to the MTBC. Our work provides compelling data to support the use of *M. riyadhense* as a novel mycobacterium for the study of virulence, evolution and pathogenesis in the MTBC.

## Materials and Methods

### Ethics Statement

The research protocol was approved by the Institutional Review Board of King Fahad Medical City (Riyadh, Saudi Arabia; #16-345) and the Institutional Biosafety and Bioethics Committee of King Abdullah University of Science and Technology (Jeddah, Saudi Arabia; #18IBEC23). All adult subjects provided informed and written consent. A parent or guardian of any child participant provided informed consent on their behalf.

### Clinical reports and bacterial strains

Eight *M. riyadhense* strains were collected in Riyadh, Saudi Arabia, between June 2011 and March 2016 (Fig 1) from patients with a positive culture for *M. riyadhense* isolated from the microbiology laboratory at the King Fahad Medical City (KFMC) in Riyadh, Saudi Arabia. The patient and sample data collected included demographic and clinical characteristics, age, sex, clinical features at presentation, presence of comorbidities, including HIV co-infection, antimicrobial susceptibility, initial and modified therapy, where applicable; and treatment outcome. Once an isolate was suspected to be an NTM, the samples were sent to the reference laboratory for full identification and antimicrobial susceptibility testing. Radiographic and pathologic data were also collected.

### Culturing, DNA isolation and sequencing of bacteria

The *M. riyadhense* strains were grown on Lowenstein Jensen (LJ) slants at 37 °C for two weeks, DNA was extracted using a phenol-chloroform protocol [76], and the quality was measured by Qubit. Twenty micrograms of high-molecular-weight (HMW) DNA from the eight *M. riyadhense* strains was sequenced using a PacBio RSII sequencer (Pacific Biosciences, Menlo Park, USA) with a 10 kb library. A NEBNext Ultra II DNA library preparation kit (New England BioLabs, Massachusetts USA) was used to prepare libraries according to the manufacturer’s instructions, and sequences from each library were generated for all *M. riyadhense* strains using the Illumina HiSeq 4000 platform (Illumina, San Diego, United States).

### Genome assembly and annotation

The Illumina short reads were trimmed, and low-quality reads were removed by Trimmomatic [77]. Eight consensus genomes based on each strain were assembled into contigs with the PacBio long reads using the Canu assembler [78]. After assembly, the draft genome was then corrected with short Illumina reads using Pilon [79] software. The circularity of assemblies was checked by Gepard [26], and assemblies were annotated by Prokka [80]. A circular map of the chromosome was compared with that of *M. tuberculosis* H37Rv and visualized with BRIG [81]. The genome of the *M. riyadhense* MR226 strain was used as a high-quality reference in this study.

### Comparison of chromosomal and plasmid gene contents of *M. riyadhense* to those of various *Mycobacterium* species

DNA sequences of 152 mycobacterial species and 77 mycobacterial plasmids were obtained from the NCBI genome database and independently annotated by Prokka [80]. The predicted protein sequences from the chromosome and each of the two plasmids (pMRLP01 and pMR01) of the *M. riyadhense* MR226 strain were then compared with the annotated genes from the rest of the mycobacterial species using Proteinortho[82]. The obtained orthologues were visualized with the heatmap package in R [83]. A focused OrthoMCL[84] comparison was performed between (1) *M. riyadhense*, *M. marinum*, *M. kansasii*, *M. szulgai* and *M. tuberculosis* and (2) *M. riyadhense* and five species from the MTBC, namely, *M. tuberculosis*, *M. bovis*, *M. canettii*, *M. mungi*, and *M. africanum*.

### SNP calling and phylogeny based on SNPs

The corrected Illumina reads were mapped using BWA[85] on to the *M. riyadhense* MR226 genome assembly. Picard tool[86] was used to clean SAM files, fix mate-pair information and mark duplicates. SNPs were called for two iterations and filtered (QD<2.0, FS>60.0, SOQ>4.0, ReadPosRankSum<-8.0) with Genome Analysis Toolkit (GATK)[87]. An alignment file was generated by SVAMP[88], and phylogeny was generated by RaxML[89] with the TVM model.

### Phylogeny of *M. riyadhense*

The PhyoPhlAn2[90] pipeline was used to identify protein sequences from 400 conserved genes in the pangenome datasets and the *Mycobacterium* genomes available at NCBI or JGI. A total of 149 species were used and *Nocardia abscessus* was used as the out-group.

A whole-genome phylogenetic tree was constructed with the MTBC species (*M. tuberculosis*, *M. bovis*, *M. canettii*, *M. mungi*, *Mycobacterium orygis*, *M. africanum*) and with *M. kansasii*, *M. marinum*, *M. shinjukuense*, *Mycobacterium leprae*, *Mycobacterium smegmatis*, *Mycobacterium parmense*, *Mycobacterium avium* and *Mycobacterium abscessus*. The one-to-one orthologues of each species were obtained using OrthoMCL and concatenated, then aligned with Muscle[91] and trimmed with TrimAL[92]. The concatenated sequences were composed of 906 genes encoding 296,124 amino acids and were used to build a phylogenetic tree with the LG+G+F model, which was selected by the ModelGenerator. The phylogenetic tree was generated by RaxML[89].

### Toxin/antitoxin systems, *mce*/*mce*-associated genes and ESX systems in *M. riyadhense* and other mycobacteria

One hundred fifty-eight T/A proteins belonging to the VapBC, RelEF, HigBA, MazEF, ParDE and UCAT families were downloaded from the NCBI protein database. We identified the *M. tuberculosis* T/A orthologues from all of the 149 species by Proteinortho[82], and the orthologue groups were also examined by Blast+ 2.4.0. The same pipeline was also applied for the MCE/MCE-associate genes, PhoPR, DosRS, PE/PPE, PE-PGRS, and ESX1-5.

### Infection of the RAW 246.7 cell line with *M. riyadhense*, *M. kansasii* and *M. bovis* BCG Denmark

The murine macrophage RAW264.7 cell line obtained from the American Type Culture Collection (ATCC, Manassas, USA) was cultured in Dulbecco’s modified Eagle’s medium (DMEM) (ThermoFisher Scientific, Waltham, USA) supplemented with 10% FCS, streptomycin, and penicillin. *M. riyadhense*, *M. kansasii* (subtype I), and *M. bovis* BCG Denmark strains were grown in Middlebrook 7H9 liquid medium after single-colony isolation from LJ slants or 7H10 agar. 7H9 was supplemented with 10% albumin, dextrose and catalase (ADC), while 7H10 was supplemented with oleic acid, albumin, dextrose and catalase (OADC) in addition to 0.2% glycerol. Ready to use LJ slants were provided by Saudi Prepared Media Laboratory (SPML, Riyadh, Saudi Arabia).

Before the infection, all bacterial cultures were centrifuged at 1,000xg for 10 minutes. The supernatant was discarded, and 10-15 3 mm glass beads were added to the pellet and then vortexed for 1 minute to break up clumps. Six ml of DMEM was added to the pellet and left to rest for 5 minutes. The upper 5 ml of the suspension was removed to a fresh 15 ml falcon tube, which was then centrifuged for an additional 3 minutes at 200xg to remove remaining bacterial clumps. The supernatant was then taken and passaged using a 26G hypodermal syringe approximately 15 times to further break up any clumps of bacteria. The optical density of the culture was measured again before the infection experiment.

RAW264.7 cells were seeded at 2 x 10^5^ cells per well in 24-well flat-bottom tissue culture plates 24 hours prior, reaching 5 x 10^5^ cells per well. The DMEM over the cells was removed, and the cells were washed once with phosphate-buffered saline (PBS). The cells were infected by applying 1 ml of DMEM containing the prepared mycobacteria from the former steps at an appropriate concentration to reach a MOI of 5:1, with DMEM alone used for the control wells. The culture plates were then returned to the incubator at 37°C with 5% CO_2_ for 3 hours to allow for bacterial uptake by the RAW264.7 cells. The supernatant was removed after 3 hours, and the infected cells were washed with PBS to remove extracellular bacteria. Subsequently, the cells were incubated in fresh DMEM with 10% FCS for 24 hours and 48 hours. For harvesting, 400 μl of TRIzol (ThermoFisher Scientific, Waltham, USA) was added to the wells at each time point, and the adherent cells were scraped out and stored at −80°C for RNA extraction. Each bacterial infection was performed in triplicate, in addition to the non-infected controls. RNA was isolated from the samples using the Direct-zol^TM^ RNA Miniprep kit (Zymo Research, Irvine, USA) according to the manufacturer’s instructions.

An Agilent RNA 6000 Nano kit was used to check the quality and quantity of the total RNA. The NanoString murine nCounter Myeloid Innate Immunity Gene Expression Panel (NanoString Technologies, USA) was used to assess transcript abundance across infections and time points using the nCounter MAX Analysis System (NanoString Technologies, Seattle, USA). The counts obtained were normalized using the nSolver^TM^ Advanced Analysis plugin (NanoString Technologies, Seattle, USA) using the geNorm algorithm, and differential gene (DE) expression was analyzed using multivariate linear regression in nSolver^TM^ software, with 0.05 as the p-value cutoff.

### Thin-layer chromatography analysis of lipooligosaccharides in *M. riyadhense*, *M. kansasii* and *M. marinum*

For TLC analysis, mycobacterial strains were grown at 30°C (*M. marinum*) or 37°C (*M. smegmatis*, *M. kansasii*, *M. riyadhense*) on LJ slants, and after sufficient incubation, grown cells were collected and washed once with PBS. Apolar and polar lipids were extracted from the cell pellets using methods described by Dobson *et al*[64]. Polar lipids were analyzed by 2D-TLC using solvent system E, which is designed to separate phospholipids and LOSs[64]. Glycolipids were visualized by charring following staining with either molybdophophoric acid (MPA) or alpha-napthol (for glycolipids).

### Diagnostic PCR markers for *M. riyadhense*

To develop diagnostic markers for *M. riyadhense* for potential use in clinical diagnosis as well as environmental and animal studies, unique regions within the *M. riyadhense* reference genome compared to that of 152 other mycobacterial species were detected using Shustring[93]. These regions were also examined by Proteinortho[82] and Blastn[94]. *mr_00036*, *mr_00263*, *mr_00606*, and *mr_01005* were selected as the amplification targets. Two primers for each gene were designed in this study:

MRDP-MR_00036-F (5’-TTCGTTGTCGGTTTCGTCGC-3’) and MRDP-MR_00036-R (5’-GCGTCAGCTCCACCGAAAAC-3’);

MRDP-MR_00263-F (5’-CCACCGCTGTTGGCGA-3’) and MRDP-MR_00263-R (5’-TTCGTCCCGTTGATCCCGTT-3’);

MRDP-MR_00606-F (5’-AACCTGCCCGATACGCACTT-3’) and MRDP-MR_00606-R (5’-ACTGTTCCTCCGTGGGGTTG-3’);

MRDP-MR_01005-F (5’-GACTGTGGGGTAACGGTGGA-3’) and MRDP-MR_01005-R (5’-CCGGTGATGTCGCCTACTCC-3’).

PCR was performed in a 25 µl reaction volume with 12.5 µl of GoTaq ® Green Master Mix (Promega, USA), 1 µl of 100 ng/µl gDNA, 1 µl with 10 nmol of forward and reverse primers, 3 µl of dimethyl sulfoxide (DMSO) and 19 µl of nuclease-free water. The PCR mixture was denatured for 5 minutes at 94°C; followed by 35 cycles of amplification involving a denaturation step at 94°C for 30 seconds, a primer annealing step at 59°C for 45 seconds, and a primer extension step at 72°C for 45 seconds; and a final extension step at 72°C for 7 minutes. The ITS-F/mycom-2 primer set[95], which is a *Mycobacterium* genus-specific primer set, was used as a control, with amplification conditions as described previously[95]. The products were electrophoresed in a 2% agarose gel for 60 minutes and visualized.

## Acknowledgements

We wish to thank members of the Biological Core Lab (BCL) of King Abdullah University of Science and Technology for sequencing the bacterial genomic DNAs on the Illumina HiSeq 4000 and PacBio RSII platforms.

## Accession codes

The *M. riyadhense* dataset is available at European Nucleotide Archive (ENA) under the study accession no. PRJEB32162. The assemblies are available with DOI 10.5281/zenodo.2873972.

## Author contribution statement

AP conceived the comparative genomics part of the study, obtained the funding and supervised the work; MG and AH conceived the clinical part of the study; MG and FA wrote the clinical part of the text; FA, MG, TA, SF, AH, MAR, AR, and TS collected the microbiological and clinical information; and AP, QG, SG, AB, and FA designed the experiments. QG performed the data analysis and prepared the initial draft of the manuscript, followed by edits from AP, SG, MG, SF, AB and CM. QG, SM, ASm and JB performed the transcriptome experiment. ASi and AB performed the TLC analysis. CN and YS provided materials for the diagnostic markers and intellectual advice. All authors have commented on various sections of the manuscript, which were finally curated and incorporated in the final version by QG and AP.

## Supporting Information

**S1 Fig. Axial enhanced CT scan image of the chest from the anonymous patient infected with *M. riyadhense* MR193.** Multifocal cavitating consolidation in both lungs predominantly involving the left upper lobe associated with ill-defined ground-glass centrilobular nodules and tree-in-bud appearance on both lungs were observed. The findings were suggestive of tuberculosis.

**S2 Fig. A comparison heatmap of predicted protein-coding gene orthologs from *M. riyadhense* MR226 and other mycobacteria. The orthologs were determined by Proteinortho**[82]**. The black and white spaces denote presence and absences of an ortholog in a given species respectively.** (A) Comparison of chromosome-encoded genes in 152 mycobacterial species. The box with red outline highlights MTBC. The red arrows indicate the genomic regions that are absent in the assembly GCA_002101845.1 while present in our MR226 assembly. (B) Comparison of the linear plasmid pMRLP01-encoded genes in 152 mycobacterial species genome assemblies. The box b with red outline highlights a region which shared orthologs with pMRLP01 in *Mycobacterium tusciae*, *Mycobacterium aromaticivorans*, *Mycobacterium llatzerense*, *Mycobacterium obuense*, *Mycobacterium novocastrense* and *Mycobacterium holsaticum*. (C) Comparison of the circular plasmid pMR01-encoded 88 genes in 152 mycobacterial species genome assemblies. The box c with red outline highlights the cluster of the pRAW-like plasmids.

**S3 Fig. Circular map of pMR01. The circles show from outside to inside (1-12) BlastN results against the pMR01 of various *Mycobacterium* plasmids. The corresponding plasmids used are listed on the right panel.** The innermost circle (13) represents pMR01 with the location of the predicted protein coding genes. Red colour indicates the location of the members of the *pe*/*ppe* gene family, blue indicates the *esx*-related genes and all the other predicted genes are colored in grey.

**S4 Fig. Multiple alignment of 8 *M. riyadhense* assemblies using progressive Mauve.** (A) The alignment of the 8 assemblies and (B) the alignment of the linear and circular plasmids from 8 strains. Circular plasmid is present in 3 strains and highlighted by boxes with black outlines.

**S5 Fig. Phylogenetic tree of *M. riyadhense* clinical isolates used in this study.** The *M. riyadhense* phylogenetic tree was constructed with SNP data from 8 datasets called by GATK pipeline by RaxML with the TVM model. The circles on each branch indicates the bootstrap values (above 95%, 1,000 replicates).

**S6 Fig. Genome alignments comparing selected Region of Differences (RDs) in mycobacteria. The Artemis Comparison Tool (ACT) was used to compare and visualize the annotated genome sequences against the chosen mycobacteria.** (A) The RD2 region of *M. riyadhense* MR226, *M. tuberculosis* H37Rv and *M. bovis* BCG Pasteur (B) The RD12 region of *M. tuberculosis*, *M. riyadhense* MR226 and *M. canettii* CIPT 140010059.

**S7 Fig. The relative distribution of the functions of the predicted protein-encoding genes were normalized by the genome size of each genome and compared to the *M. riyadhense* MR226 strain.**

The phylogenomic position of *M. riyadhense* within the *Mycobacterium* genus was determined by concatemer sequences of 906 shared single copy genes covering 296,124 amino acid positions. *Nocardia abscessus* was used as the out group. The concatenated sequences were used to build the phylogenetic tree with the LG+G+F model, which was selected by ModelGenerator. The phylogenetic tree was generated by RaxML with 1,000 iterations and the bootstrap values are shown above each branch.

**S8 Fig. Genetic locus map of the *pe-pgrs33* gene cluster from *M. marinum*, *M. kansasii*, *M. riyadhense*, *M. canettii* and *M. tuberculosis* (drawn to scale).**

The genes are shown with arrows and are colored according to the orthologs. The deletion event was highlighted with red arrows while the insertion event in green arrows.

**S9 Fig. Comparison of different gene clusters that encode type VII secretion systems in the *M. riyadhense* MR226 strain.**

The genes are shown with arrows and are colored according to the orthologs. The color codes for the figure are presented in the key. The black arrows indicate region-specific genes.

**S10 Fig. TLR and NFkB pathway responses across the three infections.**

Clustering of Nanostring data from infections with *M. riyadhense* (MR), *M. kansasii* (MK) and *M. bovis* BCG Denmark (Denm) over 3, 24 and 48 hrs showed commonality and variation in the transcriptional responses. Panel A and B show selected genes involved in the TLR (A) and NFkB (B) responses from the overall 754 gene panel used in this study.

**S11 Fig. Basic statistics of unique K-mers of *M. riyadhense* across 152 mycobacteria genome assemblies.**

**S1 Table. Comparison of *M. riyadhense* strains’ assemblies.**

**S2 Table. The SNPs analysis of different samples using MR226 as the reference.**

**S3 Table. *M. marinum* LOS locus gene orthologs shared with *M. riyadhense* strains.**

