## Supplementary Files for "Insights into ancestry and adaptive evolution of the *Mycobacterium tuberculosis* complex from analysis of the emerging pathogen *Mycobacterium riyadhense*"

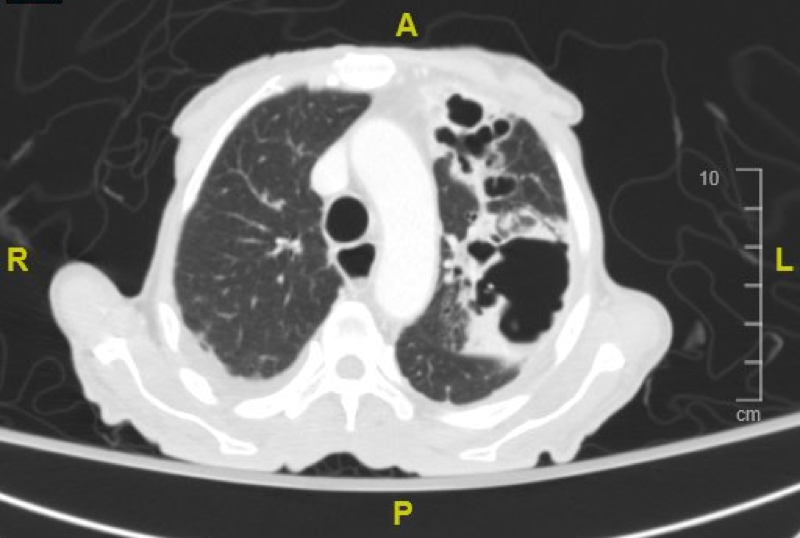

**Supplementary Fig. 1: Axial enhanced CT scan image of the chest from the anonymous patient infected with *M. riyadhense* MR193.**

Multifocal cavitating consolidation in both lungs predominantly involving the left upper lobe associated with ill-defined ground-glass centrilobular nodules and tree-in-bud appearance on both lungs were observed. The findings were suggestive of tuberculosis.

A
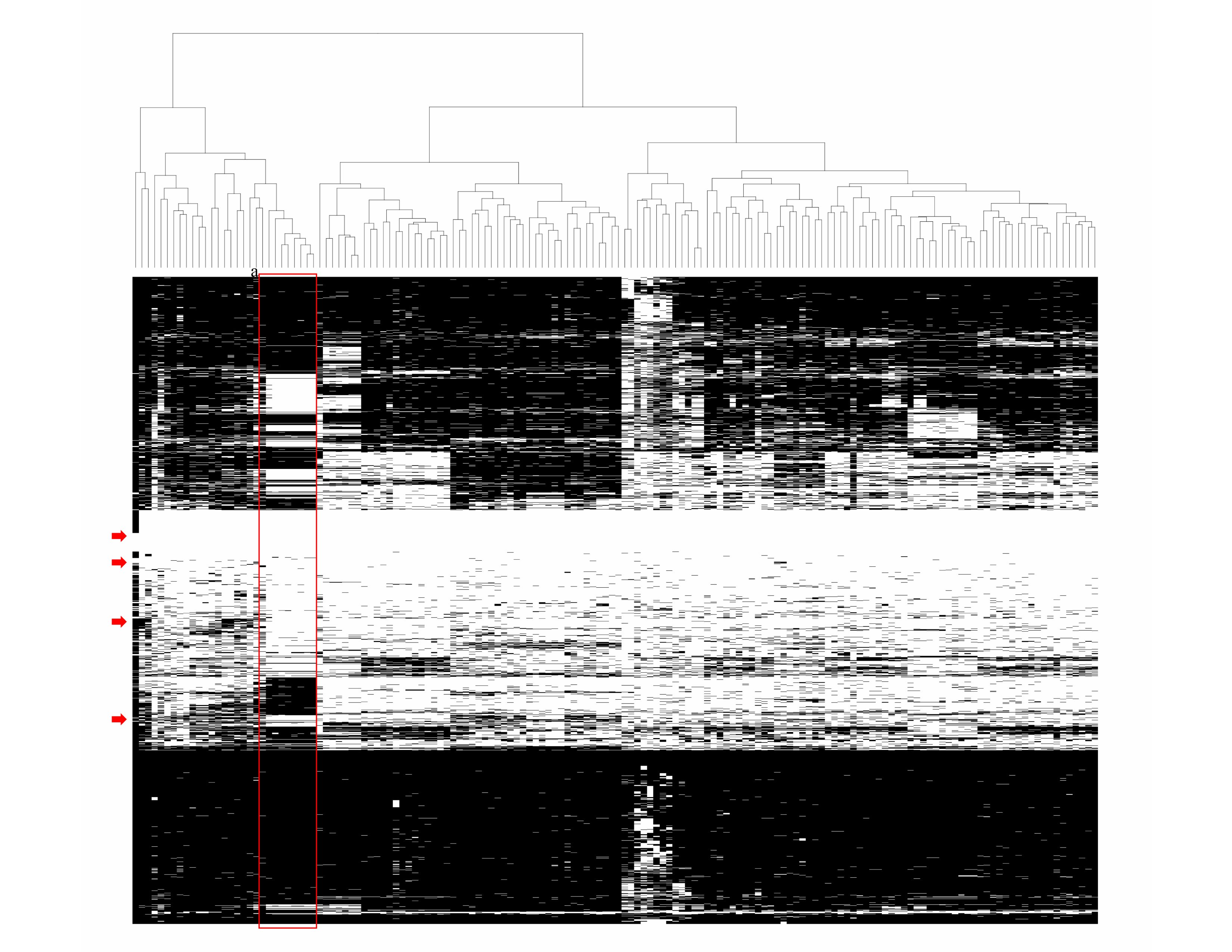

B
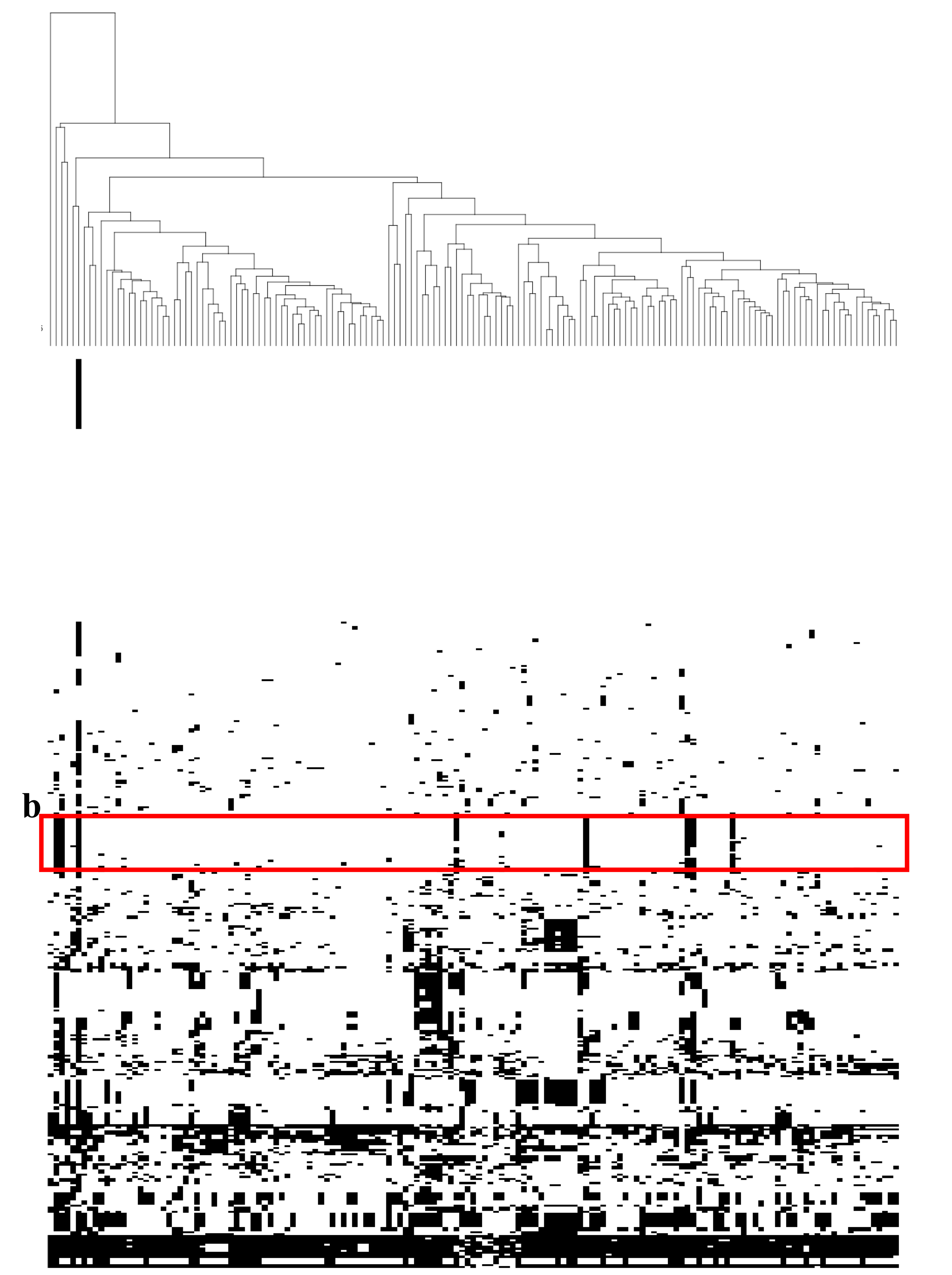

C

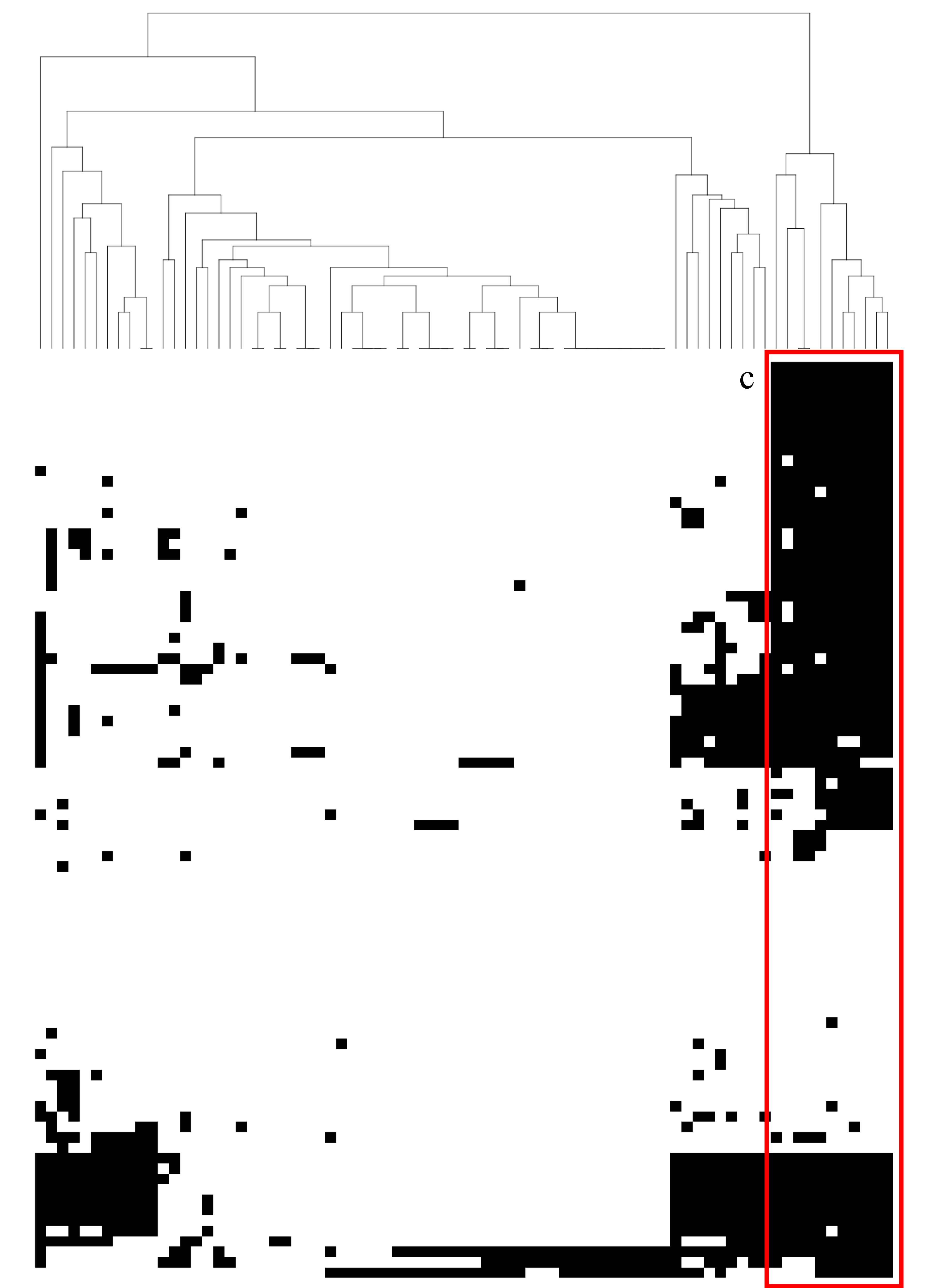

**Supplementary Fig. 2: A comparison heatmap of predicted protein-coding gene orthologs from *M. riyadhense* MR226 and other mycobacteria. The orthologs were determined by ortho-MCL. The black and white spaces denote presence and absences of an ortholog in a given species respectively.**

(A) Comparison of chromosome-encoded genes in 152 mycobacterial species. The box with red outline highlights the closely related species with *M. riyadhense* based on ortholog groups clustering. Species from left to right within box a are: *M. riyadhense* (published assembly GCA_002101845.1), *M. marinum*, *M. angelicum*, *M. tuberculosis*, *M. pseudoshottsii*, *M. liflandii*, *M. gastri*, *M. kansasii*, *M. ulcerans*, *M. shinjukuense*, *M. lacus*, *M. canettii*, *M. microti*, *M. bovis*, *M. orygis*, *M. africanum*, *M. mungi*, *M. bovis BCG.* The red arrows indicate the genomic regions that are absent in the assembly GCA_002101845.1 while present in our MR226 assembly. (B) Comparison of the linear plasmid pMRLP01-encoded genes in 152 mycobacterial species genome assemblies. The box b with red outline highlights a region which shared orthologs with pMRLP01 in *Mycobacterium tusciae*, *Mycobacterium aromaticivorans*, *Mycobacterium llatzerense*, *Mycobacterium obuense*, *Mycobacterium novocastrense* and *Mycobacterium holsaticum*; (C) Comparison of the circular plasmid pMR01-encoded 77 genes in 152 mycobacterial species genome assemblies. The box c with red outline highlights the cluster of the pRAW-like plasmids.

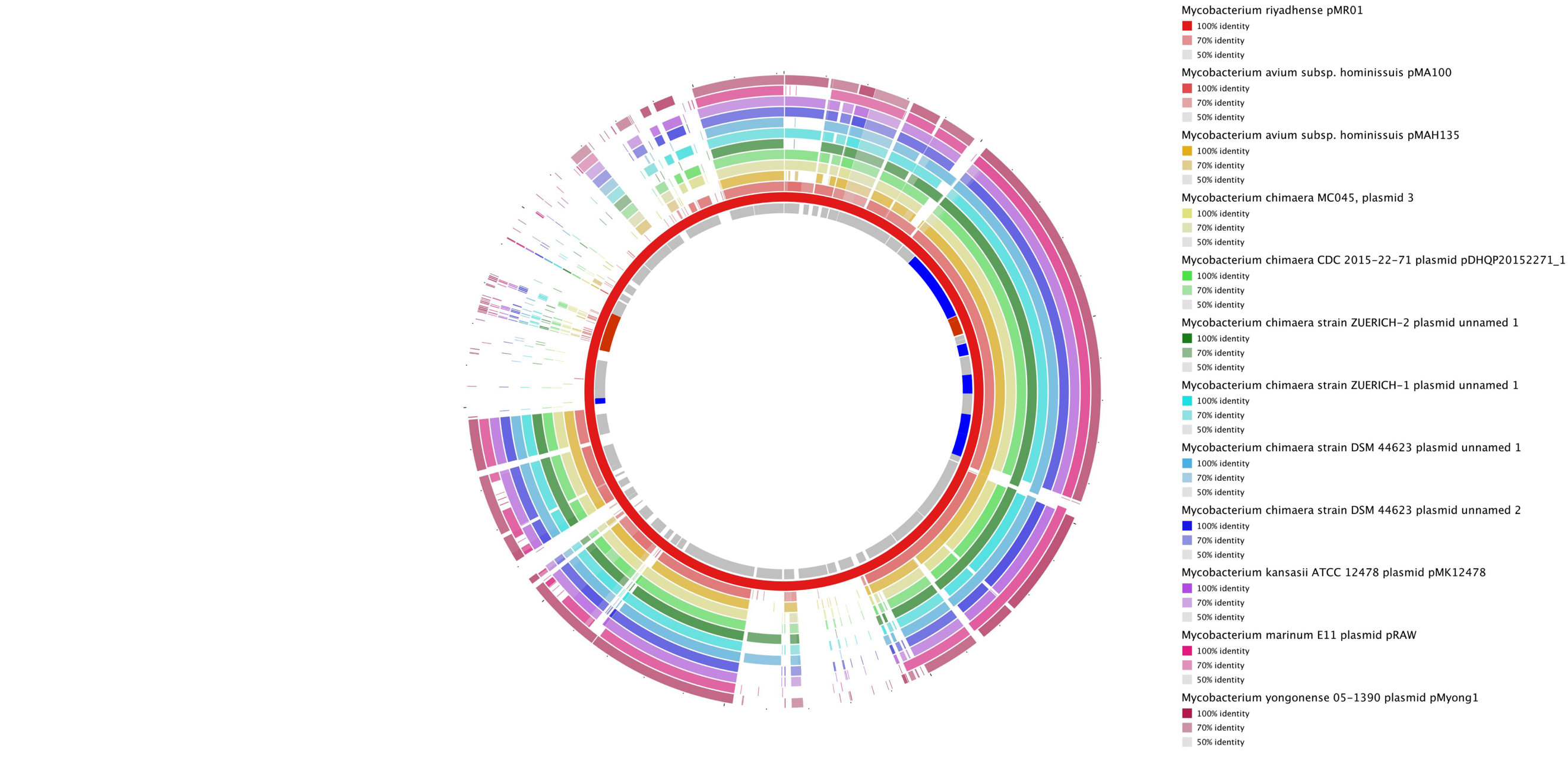

**Supplementary Fig. 3: Circular map of pMR01. The circles show from outside to inside (1-12) BlastN results against the pMR01 of various *Mycobacterium* plasmids. The corresponding plasmids used are listed on the right panel.**

The innermost circle (13) represents pMR01 with the location of the predicted protein coding genes. Red colour indicates the location of the members of the *pe*/*ppe* gene family, blue indicates the *esx*-related genes and all the other predicted genes are colored in grey. ­­­­­

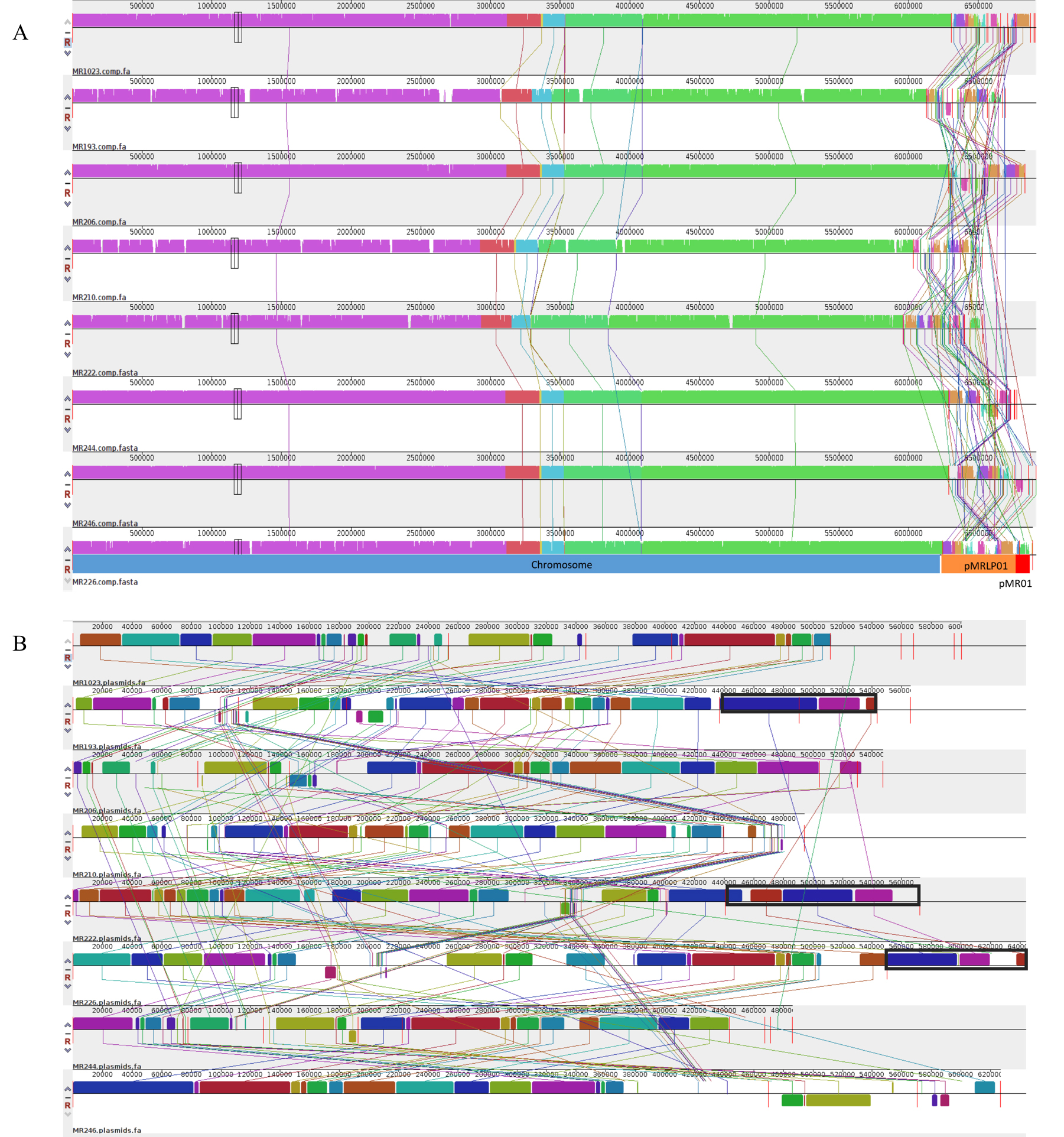

**Supplementary Fig. 4: Multiple alignment of 8 *M. riyadhense* assemblies using progressive Mauve.**

(A) The alignment of the 8 assemblies and (B) the alignment of the linear and circular plasmids from 8 strains. Circular plasmid is present in 3 strains and highlighted by boxes with black outlines.

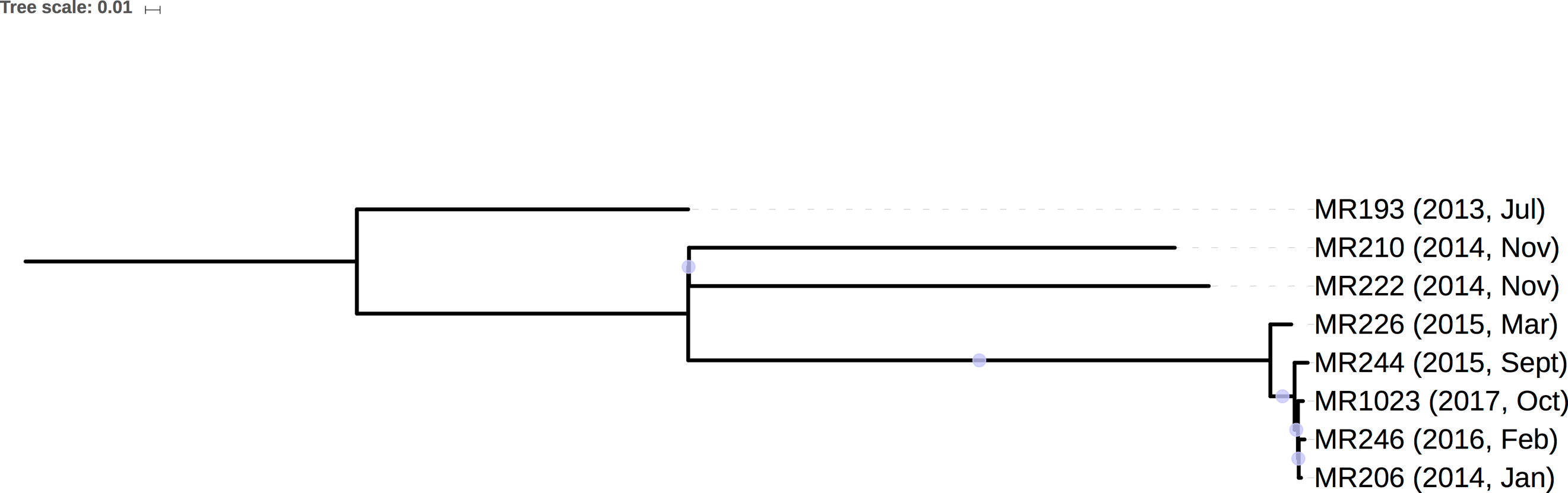

**Supplementary Fig. 5: Phylogenetic tree of *M. riyadhense* clinical isolates used in this study.**

The *M. riyadhense* phylogenetic tree was constructed with SNP data from 8 datasets called by GATK pipeline by RaxML with the TVM model. The circles on each branch indicates the bootstrap values (above 80%, 1,000 replicates).

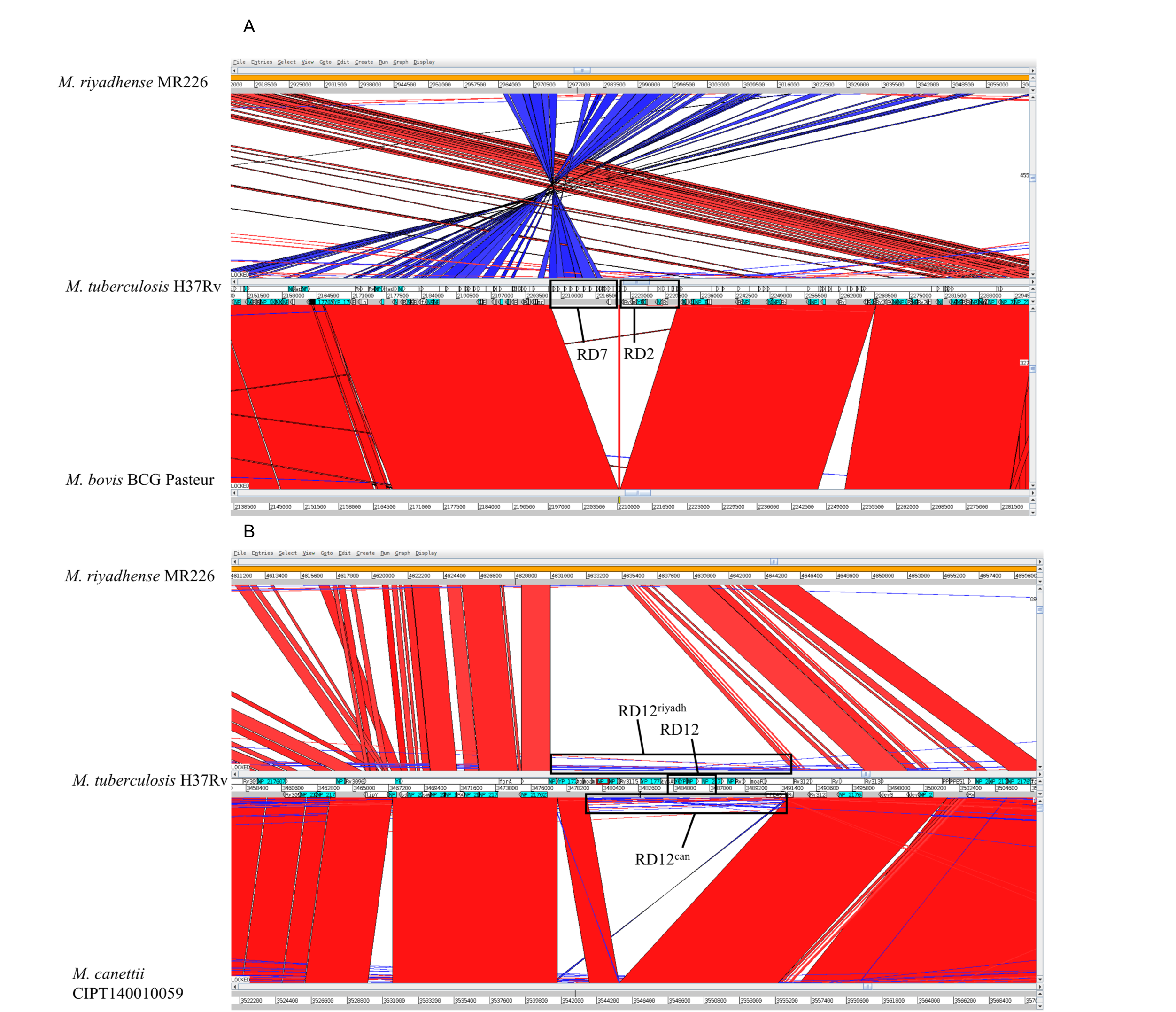

**Supplementary Fig. 6: Genome alignments comparing selected Region of Differences (RDs) in mycobacteria. The Artemis Comparison Tool (ACT) was used to compare and visualize the annotated genome sequences against the chosen mycobacteria.**

(A) The RD2 region of *M. riyadhense* MR226, *M. tuberculosis* H37Rv and *M. bovis* BCG Pasteur (B) The RD12 region of *M. tuberculosis*, *M. riyadhense* MR226 and *M. canettii* CIPT 140010059.

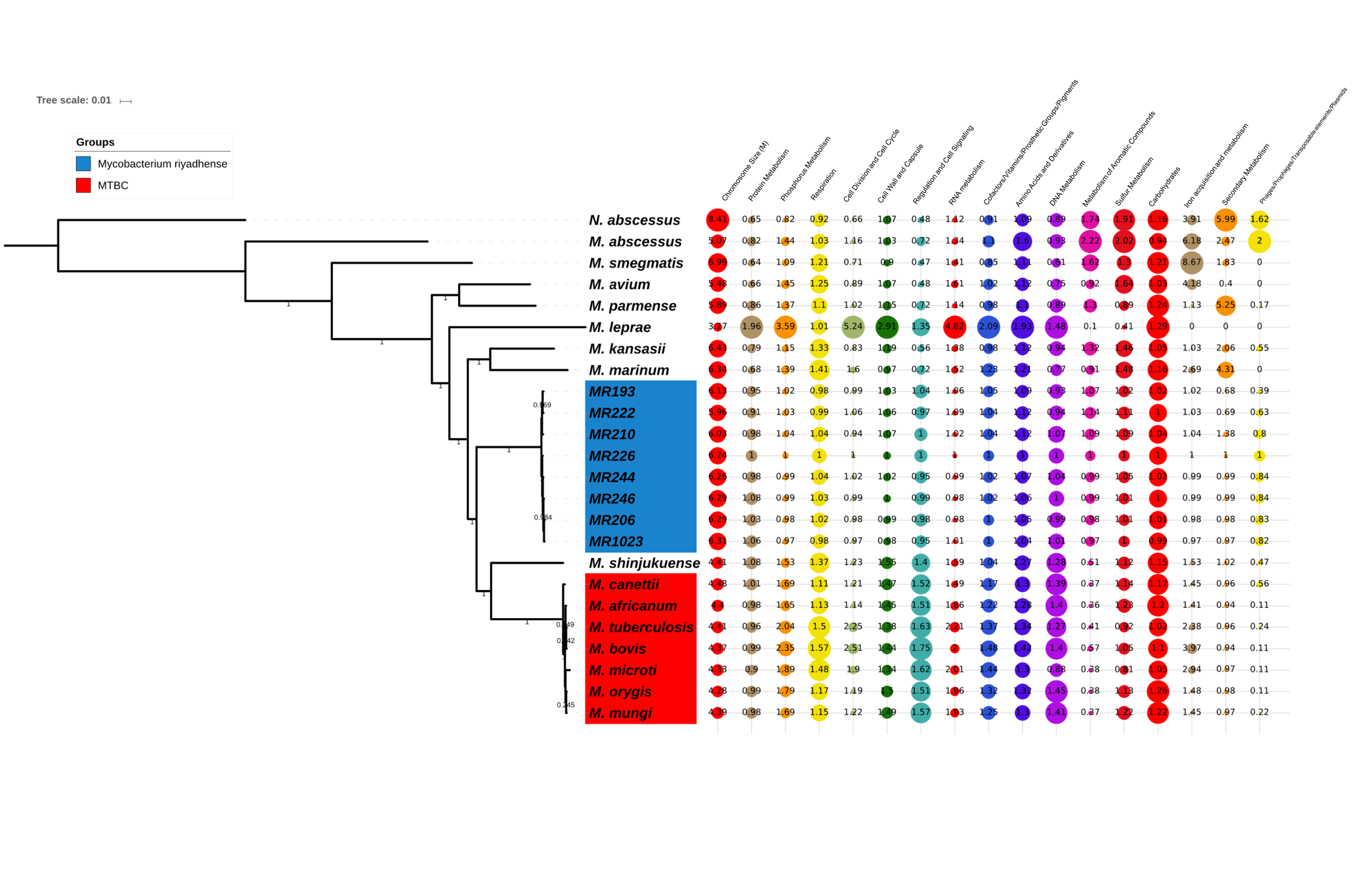

**Supplementary Fig. 7: The relative distribution of the functions of the predicted protein-encoding genes were normalized by the genome size of each genome and compared to the *M. riyadhense* MR226 strain.**

The phylogenomic position of *M. riyadhense* within the *Mycobacterium* genus was determined by concatemer sequences of 906 shared single copy genes covering 296,124 amino acid positions. *Nocardia abscessus* was used as the out group. The concatenated sequences were used to build the phylogenetic tree with the LG+G+F model, which was selected by ModelGenerator. The phylogenetic tree was generated by RaxML with 1,000 iterations and the bootstrap values are shown above each branch.

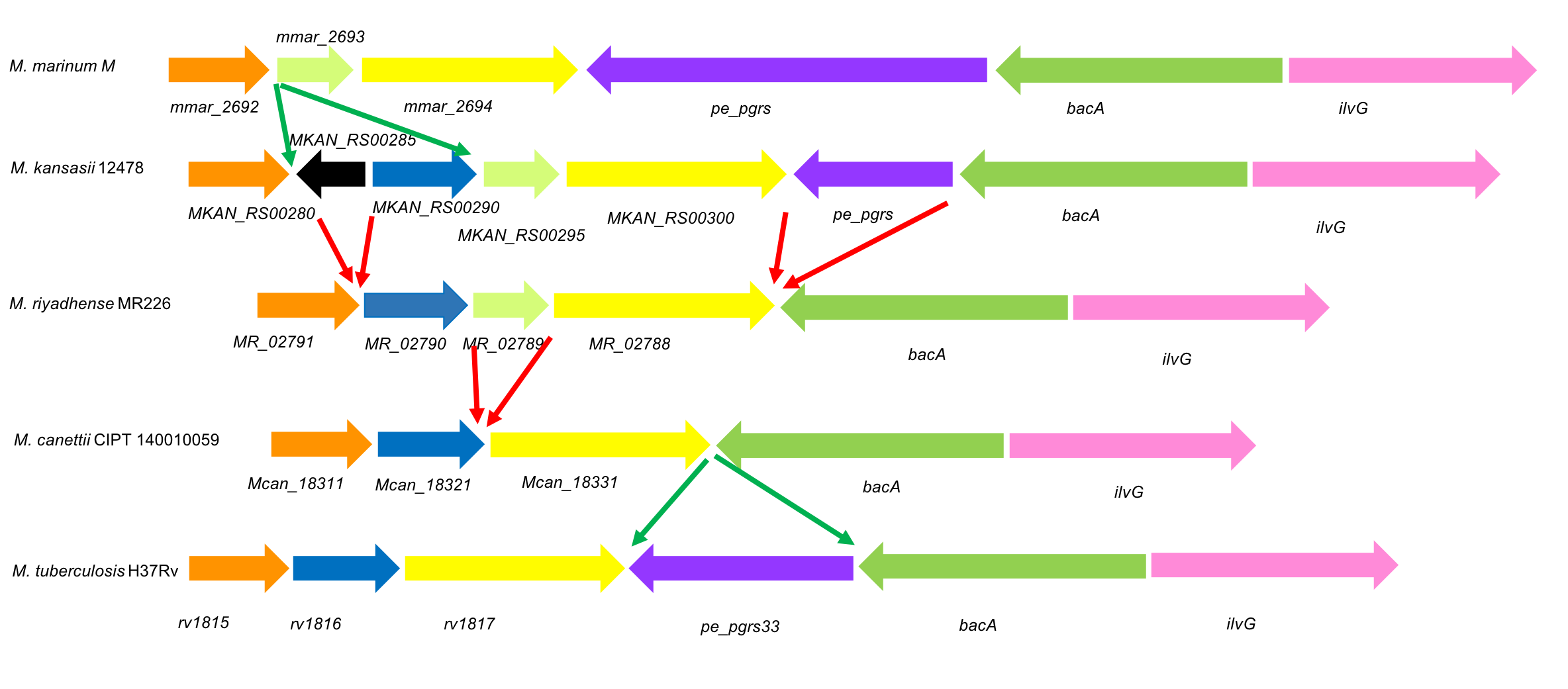

**Supplementary Fig. 8: Genetic locus map of the *pe-pgrs33* gene cluster from *M. marinum*, *M. kansasii*, *M. riyadhense*, *M. canettii* and *M. tuberculosis* (drawn to scale).**

The genes are shown with arrows and are colored according to the orthologs. The deletion event was highlighted with red arrows while the insertion event in green arrows.

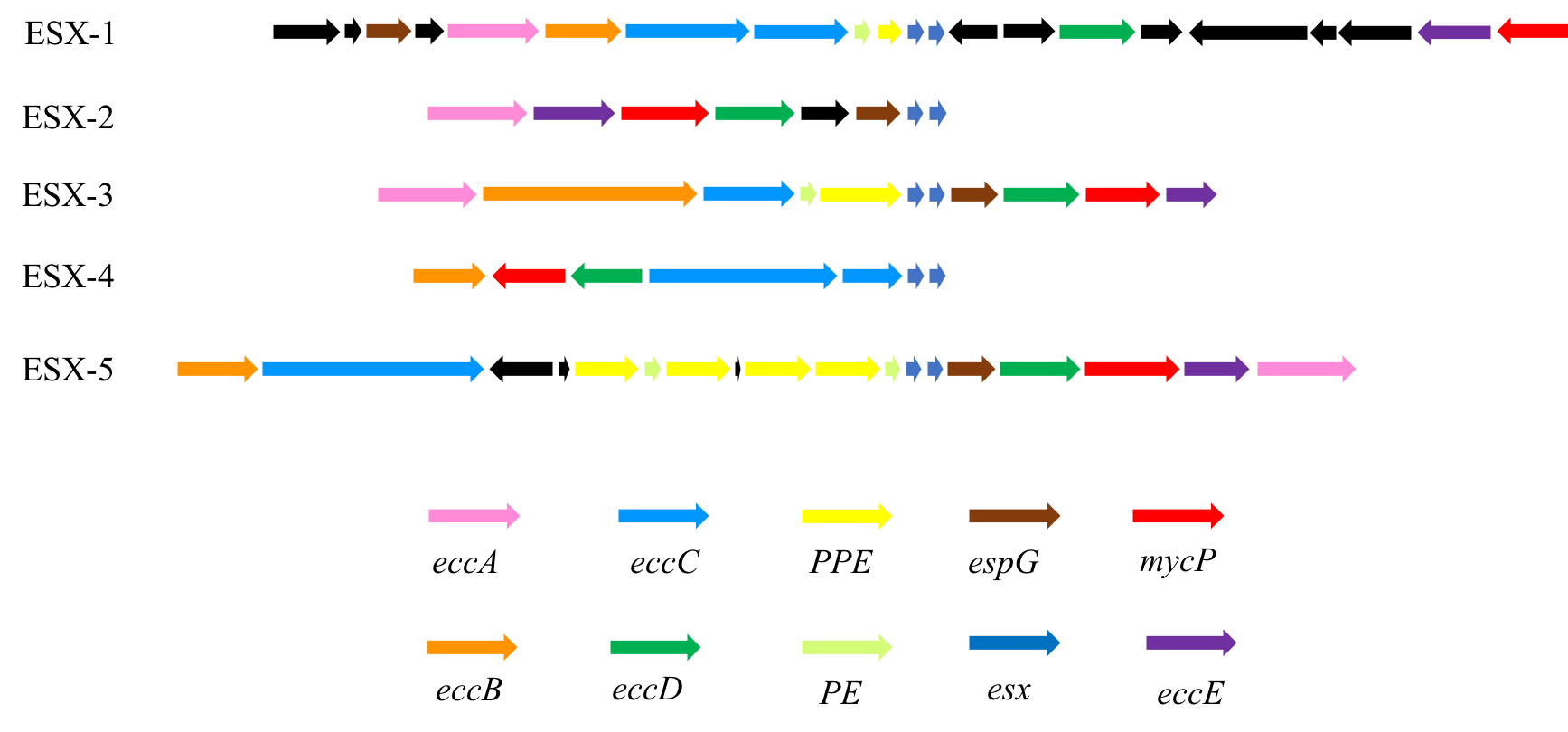

**Supplementary Fig. 9: Comparison of different gene clusters that encode type VII secretion systems in the *M. riyadhense* MR226 strain.**

The genes are shown with arrows and are colored according to the orthologs. The color codes for the figure are presented in the key. The black arrows indicate region-specific genes.

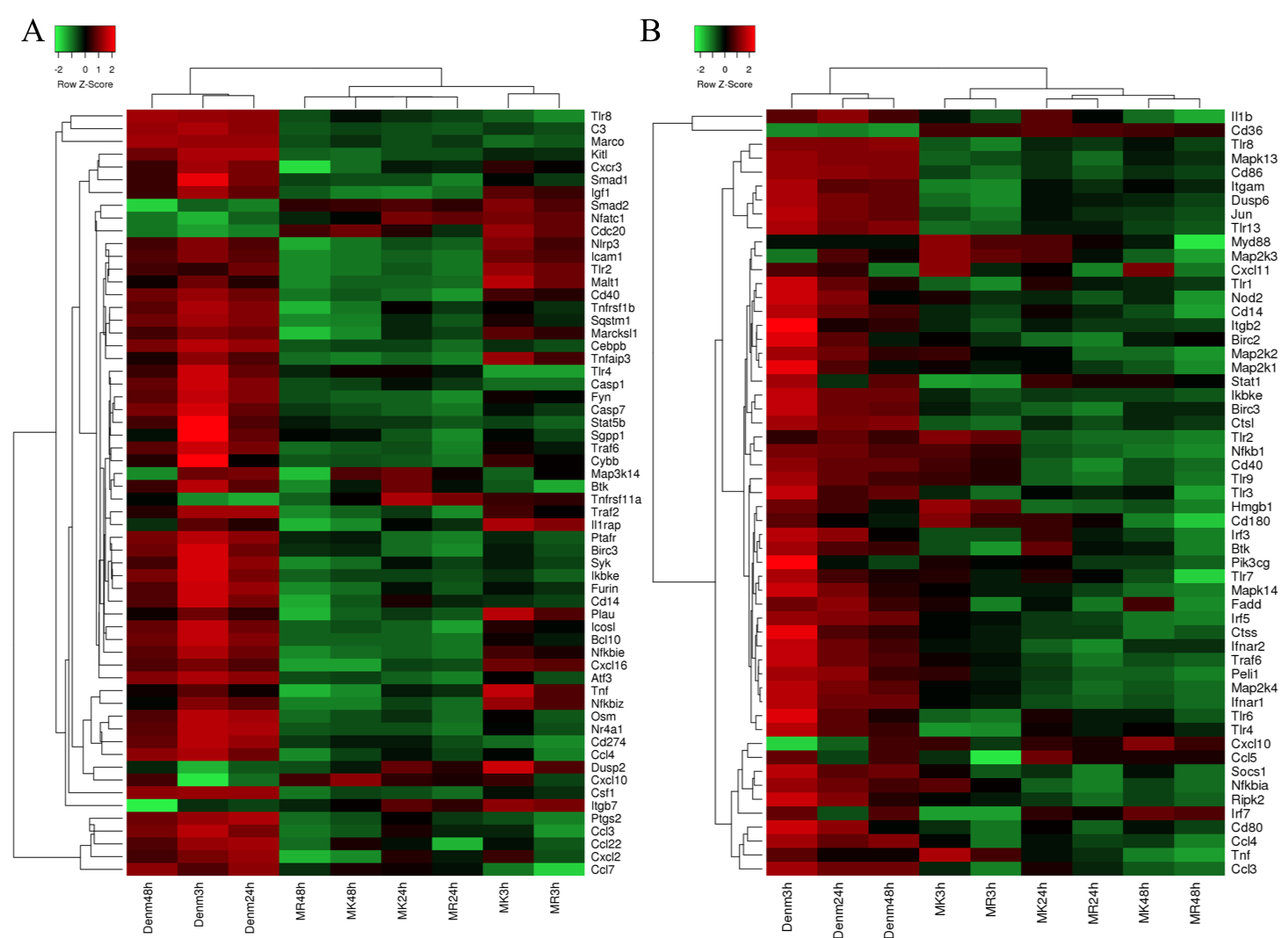

**Supplementary Fig. 10: TLR and NFkB pathway responses across the three infections.**

Clustering of Nanostring data from infections with *M. riyadhense* (MR), *M. kansasii* (MK) and *M. bovis* BCG Denmark (Denm) over 3, 24 and 48 hrs showed commonality and variation in the transcriptional responses. Panel A and B show selected genes involved in the TLR (A) and NFkB (B) responses from the overall 754 gene panel used in this study.

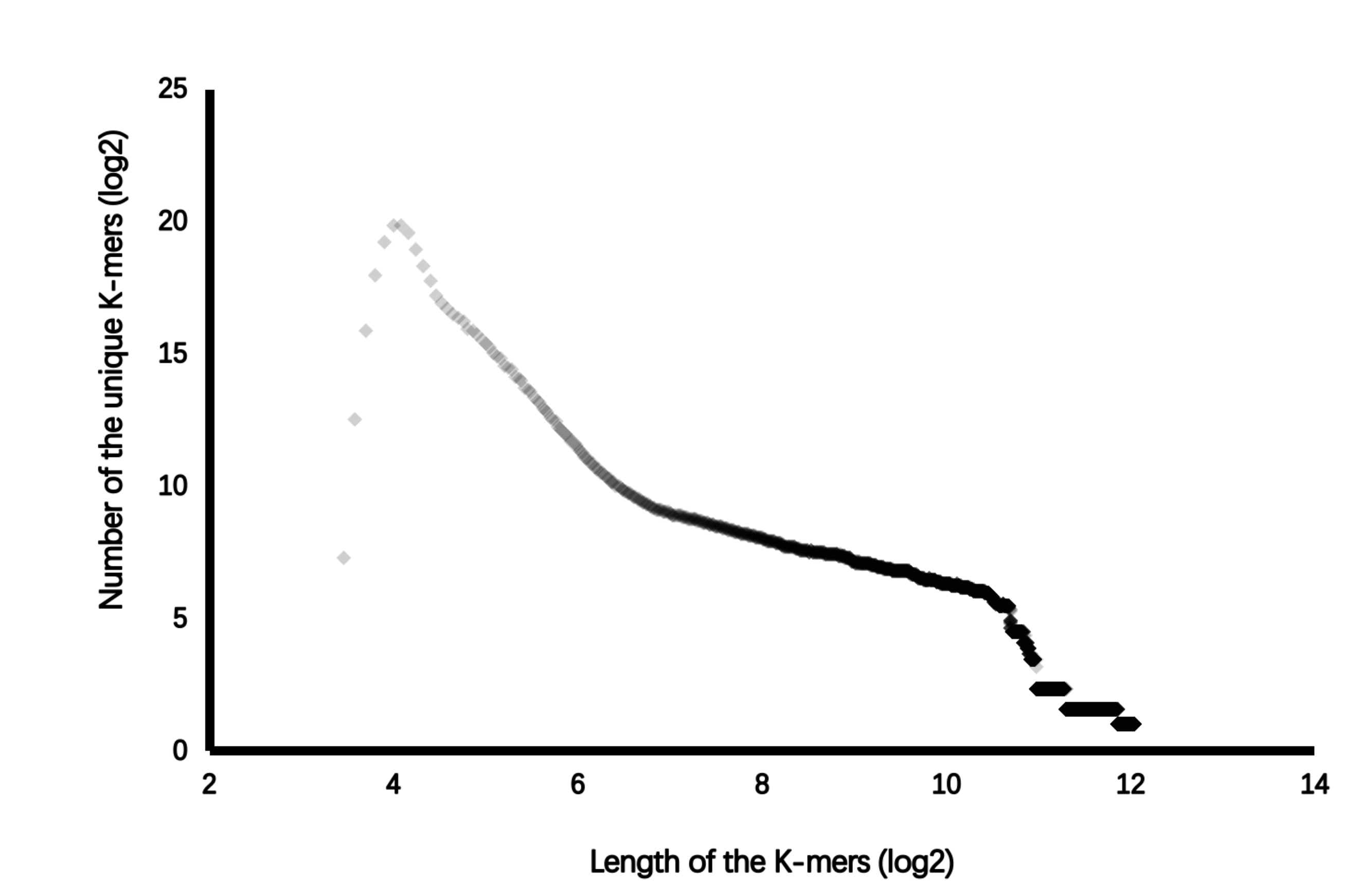

**Supplementary Fig. 11: Basic statistics of Unique K-mers of *M. riyadhense* across 152 mycobacteria genome assemblies.**

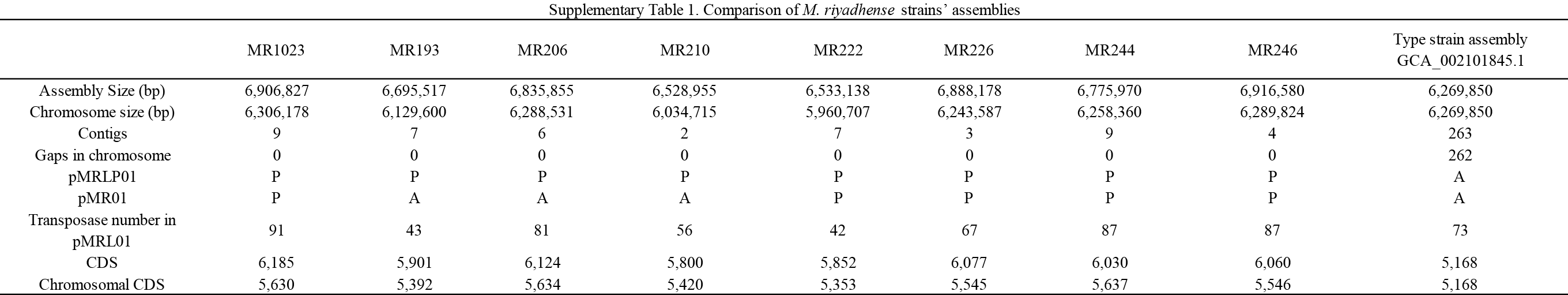

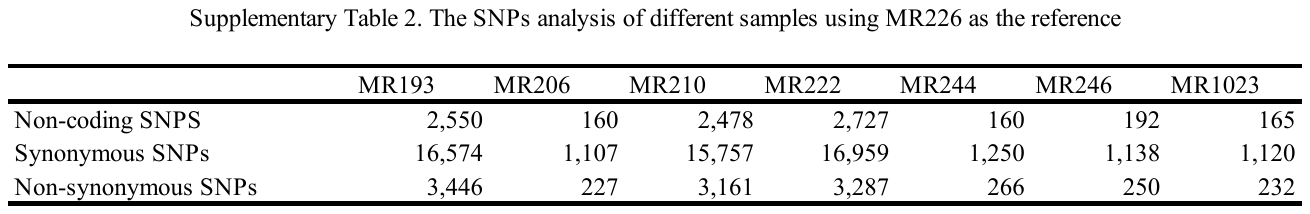

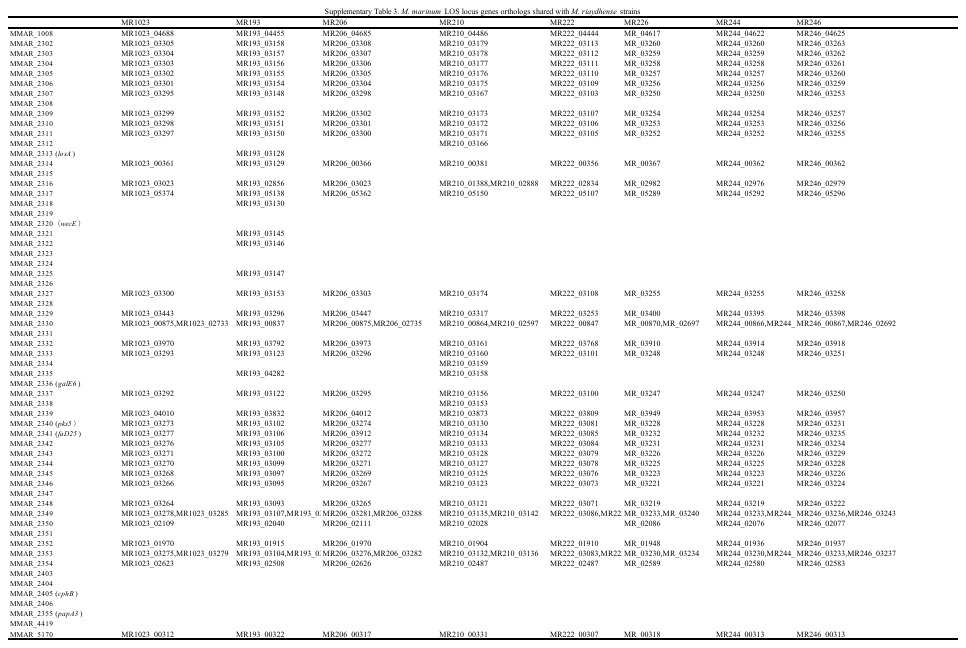
